# A Non-Canonical Function of Arabidopsis ERECTA Proteins in Gibberellin Signaling

**DOI:** 10.1101/2021.12.02.470991

**Authors:** Elzbieta Sarnowska, Szymon Kubala, Pawel Cwiek, Sebastian Sacharowski, Paulina Oksinska, Jaroslaw Steciuk, Magdalena Zaborowska, Jakub M. Szurmak, Roman Dubianski, Anna Maassen, Malgorzata Stachowiak, Bruno Huettel, Monika Ciesla, Klaudia Kogut, Anna T. Rolicka, Saleh Alseekh, Ernest Bucior, Rainer Franzen, Anna Klepacz, Malgorzata A. Domagalska, Samija Amar, Janusz A. Siedlecki, Alisdair R. Fernie, Seth J. Davis, Tomasz J. Sarnowski

**Affiliations:** Maria Sklodowska-Curie National Research Institute of Oncology, Roentgena 5, Warsaw, Poland; Institute of Biochemistry and Biophysics Polish Academy of Sciences, Pawinskiego 5A Warsaw, Poland; Max Planck Genome Centre Cologne, D-50820 Cologne, Germany; Faculty of Biology, University of Warsaw, Warsaw, Poland; Max Planck Institute of Molecular Plant Physiology, 14476 Potsdam-Golm, Germany; Center for Plant Systems Biology and Biotechnology, 4000 Plovdiv, Bulgaria; Max-Planck Institute for Plant Breeding Research; D-50829 Cologne, Germany; State Key Laboratory of Crop Stress Biology, School of Life Sciences, Henan University, 13 Kaifeng 475004, China; Department of Biology, University of York, York YO10 5DD, UK

**Keywords:** Arabidopsis, ERECTA, ERECTA-LIKE1, ERECTA-LIKE2, LRR-RLK, SWI/SNF, SWI3B, HER2, Chromatin

## Abstract

The Arabidopsis ERECTA family (ERf) of leucine-rich repeat receptor-like kinases (LRR-RLKs), comprising ERECTA (ER), ERECTA-LIKE 1 (ERL1) and ERECTA-LIKE 2 (ERL2), control epidermal patterning, inflorescence architecture, stomata development, and hormonal signaling. Here we show that the *er/erl1/erl2* triple mutant exhibits impaired gibberellin (GA) biosynthesis and perception alongside broad transcriptional changes. ERf proteins interact in the nucleus, *via* kinase domains, with the SWI3B subunit of the SWI/SNF chromatin remodeling complex (CRCs). The *er/erl1/erl2* triple mutant exhibits reduced SWI3B protein level and affected nucleosomal chromatin structure. The ER kinase phosphorylates SWI3B *in vitro*, and the inactivation of all ERf proteins leads to the decreased phosphorylation of SWI3B protein *in vivo*. Correlation between DELLA overaccumulation and SWI3B proteasomal degradation together with the physical interaction of SWI3B with DELLA proteins explain the lack of RGA accumulation in the GA- and SWI3B-deficient *erf* mutant plants. Co-localization of ER and SWI3B on *GID1* (*GIBBERELLIN INSENSITIVE DWARF 1*) DELLA target gene promoter regions and abolished SWI3B binding to *GID1* promoters in *er/erl1/erl2* plants supports the conclusion that ERf-SWI/SNF CRC interaction is important for transcriptional control of GA receptors. Thus, the involvement of ERf proteins in transcriptional control of gene expression, and observed similar features for human HER2 (Epidermal Growth Family Receptor-member), indicate an exciting target for further studies of evolutionarily conserved non-canonical functions of eukaryotic membrane receptors.

**ONE SENTENCE SUMMARY:** ERECTA leucine-rich receptor-like kinase and SWI3B subunit of SWI/SNF chromatin remodeling complex cooperate in direct transcriptional control of *GID1* genes in Arabidopsis.

## Introduction

The ERECTA family (ERf) of leucine-rich-repeat receptor-like kinases (LRR-RLKs) consists of three members: ERECTA (ER), ERECTA-LIKE 1 (ERL1), and ERECTA-LIKE 2 (ERL2). ERf proteins carry extra-cellular leucine-rich repeats (LRRs), as well as transmembrane and cytosolic kinase domains (Shpak et al., 2004; Torii et al., 1996, Kosentka et al., 2017). Inactivation of *ERECTA* leads to inflorescence, pedicels, and siliques compaction, while the individual loss of either *ERL1* or *ERL2* function has a limited effect on Arabidopsis development (Shpak et al., 2004). ERf proteins are functionally redundant-their simultaneous inactivation results in dramatic growth retardation, severe dwarfism, enlargement of the shoot apical meristem (SAM), clustered stomata, and sterility. ERf regulates stem cell homeostasis *via* buffering cytokinin responsiveness and auxin perception in SAM and modulating the balance between stem cell proliferation and consumption (Shpak et al., 2004; Griffiths et al., 2006; Torii et al., 2007; Chen et al., 2013; Shpak, 2013; Uchida et al., 2013; Zhang et al., 2021). ERECTA controls the expression of genes associated with gibberellin (GA) metabolism (Uchida et al., 2012a) restricting xylem expansion downstream of the GA pathway (Ragni et al., 2011). It additionally regulates shade avoidance in a GA and auxin-dependent manner (Du et al., 2018) and ethylene-induced hyponastic growth (Van Zanten et al., 2010).

Overexpression of ER variant lacking the C-terminal kinase domain (ERΔK) caused more severe developmental defects than complete inactivation of *ERECTA,* suggesting an interaction of the kinase domain with important regulatory partners (Shpak, 2003). ERECTA interacts with ERL1 and ERL2 to form receptor complexes recognizing two endodermis-derived peptide hormones (EPFL4 and EPFL6), regulating vascular differentiation and stem elongation. ERf proteins additionally form complexes with the receptor-like protein TOO MANY MOUTHS (TMM), which controls stomatal differentiation by recognition of the secretory peptides EPIDERMAL PATTERNING FACTOR 1 (EPF1), EPF2, and stomagen (Lee et al., 2012; Uchida et al., 2012a; Lee et al., 2015b).

ERL2 has been found to undergo endocytosis (Ho et al., 2016), suggesting that ERf proteins may play, as yet uncharacterized, regulatory roles upon internalization, in addition to their functions as ligand-binding membrane receptors. ERECTA signaling, in tandem with the SWR1 chromatin remodeling complex (CRC), controls the expression of the *PACLOBUTRAZOL RESISTANCE 1* (*PRE1*) family genes. This observation supports their role in the GA signaling pathway, however, neither direct interaction between ERECTA and SWR1 nor the direct influence of ERECTA signaling on chromatin structure or SWR1 activity has, as yet, been demonstrated (Cai et al., 2017; Cai et al., 2021).

Here we show that the loss of all ERf proteins in the *er/erl1/erl2* triple mutant (*erf*) results in broad transcriptomic changes affecting hormonal, developmental, and metabolic processes. Inactivation of ERf proteins caused down-regulation of the GA receptor *GID1* (*GIBBERELLIN INSENSITIVE DWARF 1*) genes expression and decreased bioactive GA levels. The ER protein undergoes endocytosis and enters the nucleus. All three ERf proteins interact in the nucleus with the SWI3B core subunit of the SWI/SNF CRCs. The kinase domain of the ER protein exhibits the ability to phosphorylate SWI3B protein. The physical interaction of SWI3B with RGA and RGL1, together with identified correlation between DELLA accumulation and SWI3B proteasomal degradation, provide an explanation as to why GA-deficient *erf* mutant plants did not overaccumulate RGA. These data collectively suggest cooperation of ERf-signaling with SWI/SNF in the modulation of gene transcription. The ER and SWI3B also co-localized in the promoter regions of *GID1* DELLA target genes. In the *erf* mutant, the binding of SWI3B to *GID1* promoters was abolished. These results collectively suggest that ERf proteins directly control GA receptor expression by restricting recruitment of the SWI/SNF CRCs to its target *loci*.

## Results

### Inactivation of *Erf* Proteins Has a Broad Effect on the Arabidopsis Transcriptome including GA Signaling

The Arabidopsis *er/erl1/erl2* plants exhibit severe dwarfism, dark green color, defects in vascular development, stem elongation, and stomatal differentiation, as well as complete sterility (Figure 1 A, Supplemental Figure 1A, (Shpak et al., 2004)).

**Figure 1.**
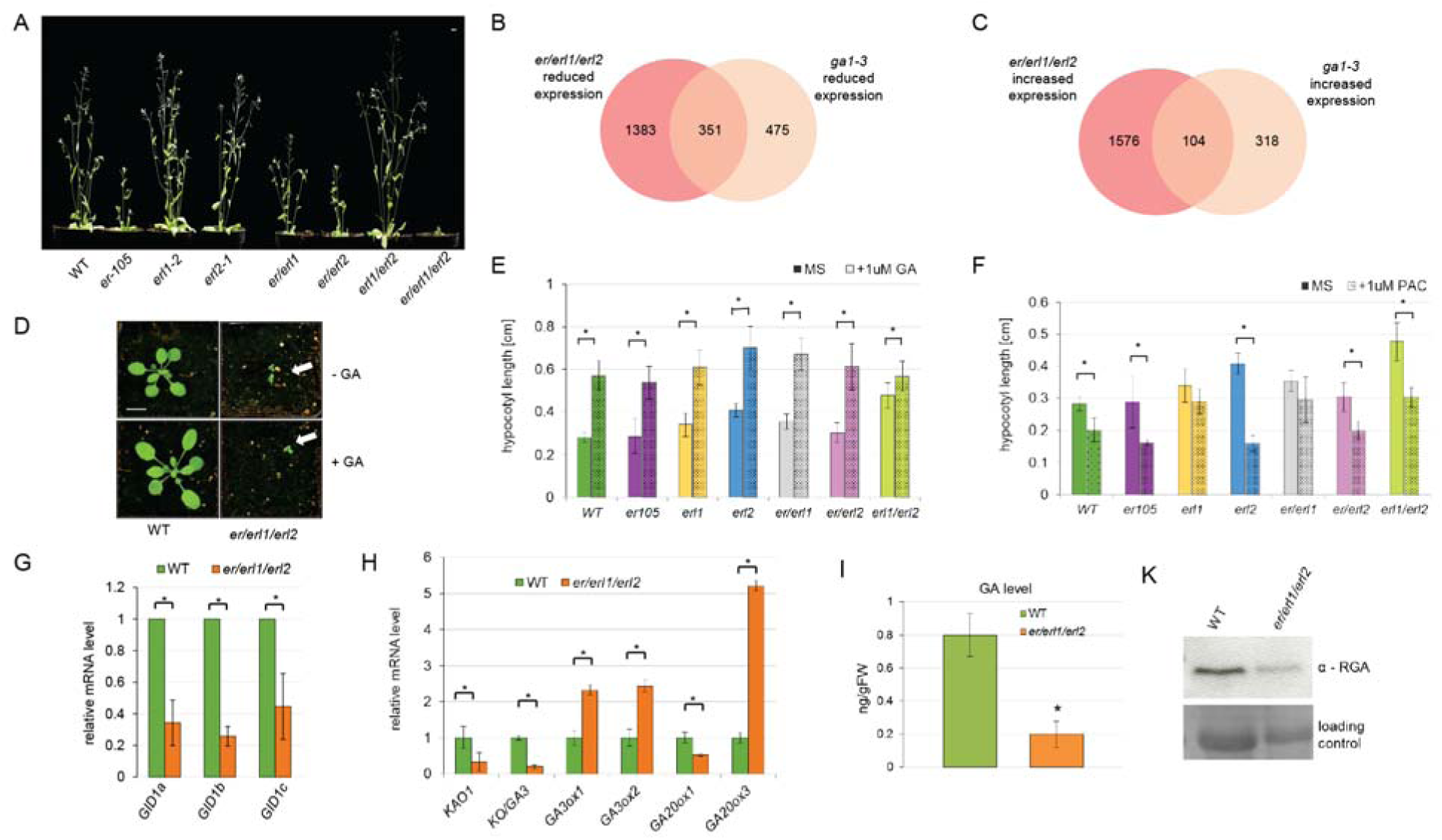
*ERf* inactivation affects Arabidopsis development, causes transcriptomic changes overlapping with the effect of *ga1-3* mutation and impairs GA biosynthesis and signaling (See also Figures S1, S2 and S3). A, Phenotypic changes conferred by combinations of *erf* mutations. Scale bar= 1 cm. B, Overlapping down-regulated genes in *er/erl1/erl2* and *ga1-3* plants. C, Overlapping up-regulated genes in *er/erl1/erl2* and *ga1-3* plants. D, The *er/erl1/erl2* plants exhibit impaired GA response. 14-days old LD (12h day/12 night) grown WT and *er/erl1/erl2*, sprayed twice a week with water (upper row) or 100µM GA_4+7_ (lower row). Arrows-*er/erl1/erl2* plants. Scale bar= 1cm. E, The GA response is retained to various levels in combinations of *erf* mutants. Error bars-SD,* = P < 0.05, Student’s *t-*test, *n*= 30 plants. F, The response of various *erf* mutants to 1µM Paclobutrazol treatment. Error bars-SD,* = P < 0.05, Student’s t test, n= 30 plants. G, The *er/erl1/erl2* mutant exhibits altered transcription of *GID1* GA receptor genes (error bars-SD, P < 0.05, Student’s *t*-test, three biological and technical replicates were assayed). H, The *er/erl1/erl2* mutant displays altered GA biosynthesis and metabolism-related genes expression (error bars-SD, P < 0.05, Student’s *t*-test, three biological and technical replicates were assayed). I, The *er/erl1/erl2* mutant exhibits dramatically reduced level of bioactive GA_4+7_ gibberellin (error bars-SD, P < 0.05, Student’s *t*-test, three biological and technical replicates were assayed). J, The *er/erl1/erl2* mutant shows decreased level of the DELLA protein RGA.

Given the severe phenotypic alterations of the *er/erl1/erl2* plants, we performed transcript profiling with Affymetrix ATH1 microarrays on RNA samples from aerial parts of the *er/erl1/erl2* mutant and WT (wild type) adult plants (representing the most comparable stage of the development) grown for 5 weeks under long-day conditions (16h day/8h night). Data analysis identified 1734 versus 1680 genes showing >1.50-fold decrease and increase, respectively, of transcript levels in *er/erl1/erl2* comparing to WT (Supplemental Figure 1B, Supplemental Dataset 1 Sub-tables 1, 2). Gene Ontology (GO) terms of primary metabolism, developmental processes, and response to hormones were enriched among the *er/erl1/erl2* down-regulated genes (Supplemental Dataset 1 Sub-table 3). Among these, 27 genes were classified to GA-response (Supplemental Table 1, Supplemental Dataset 1 Sub-tables 1, 3). The up-regulated genes were classified into GO-terms of chloroplast-related metabolic and light-regulated transcription processes, responses to cytokinin, and auxin degradation (Supplemental Dataset 1 Sub-Tables 2,4). Several genes acting in leaf epidermal and stomatal cell differentiation showed enhanced transcription in the *er/erl1/erl2* mutant (Supplemental Table 2). In conclusion, the inactivation of ERf altered transcriptional regulation of hundreds of targets, including a set of GA-regulated genes.

Phenotypic traits exhibited by double and triple *erf* mutants resemble those of double and triple *gid1abc* (*gibberellin insensitive dwarf 1a, b* and *c*) plants (Figure 1A; (Griffiths et al., 2006)). Inactivation of *GID1abc* genes has a nearly identical effect on the Arabidopsis transcriptome as the severe GA-deficient mutant *ga1-3* (Willige et al., 2007), thus we compared the transcriptomic data available for the *ga1-3* mutant with those caused by inactivation of all *ERf* genes.

We identified a large overlap of differentially expressed genes (DEG) in the *er/erl1/erl2* and *ga1-3* lines. Among 826 genes down-regulated in the *ga1-3* line, 351 (about 42.5%) also exhibited decreased expression in the *er/erl1/erl2* plants (Figure 1B), while 104 genes (about 24.6% of *ga1-3* up-regulated genes) were up-regulated in both lines (Figure 1C). Only 33 genes were up-regulated in *ga1-3* but down-regulated in *er/erl1/erl2* (Supplemental Figure 1C), and only 64 genes down-regulated in *ga1-3* but up-regulated in *er/erl1/erl2* (Supplemental Figure 1D). DEG common to *ga1-3* and *er/erl1/erl2* lines belonged to both DELLA (repressors of GA pathway) -dependent and DELLA-independent classes (Cao et al., 2006), regardless of whether they display co-regulation or contrasting regulation in these lines (Supplemental Figure 1E and F). This suggests the involvement of Arabidopsis ERf proteins in the control of GA-related processes. Therefore, we next tested the response of *er/erl1/erl2* plants to exogenously supplied bioactive 100 µM GA_4+7_ and found that spraying of the *er/erl1/erl2* mutant grown under LD condition (12h day/12h night) did not lead to increased leaf size by day 14 compared to the remarkable expansion of control WT rosette leaves (Figure 1D, Supplemental Figure 2A). Thus *er/erl1/erl2* displayed GA insensitivity. Nonetheless, the GA-treatment resulted in bolting of *er/erl1/erl2* plants, only after over two months (Supplemental Figure 2B and C), indicating their residual response to GA.

Although we showed that ERf proteins are involved in the GA response, it remained unclear whether proper GA perception requires all ERf proteins. Thus, we tested the hypocotyl response of single and double *erf* mutants in various combinations to the treatment with 1 µM GA_4+7_ or 1 µM Paclobutrazol (PAC), an inhibitor of GA biosynthesis. The GA response was retained to various levels in all tested mutants (Figure 1E) while the *erl1* and *er/erl1* plants had an impaired response to PAC and *erl1/erl2* displayed a significant reduction of hypocotyl length (Figure 1F).

Upon crossing *er*, *er/erl1*, and *er/erl2* lines with the *ga1-3* mutant, we observed only a discrete enhancement of the *ga1-3* phenotype. However most of the phenotypic changes characteristic for *ga1-3* mutation were retained, indicating that many of the *er, erl1,* or *erl2* single or double mutant phenotypes are likely not exclusively a result of GA deficiency (Supplemental Figure 2D and E).

We have proven that only parallel inactivation of all ERf proteins causes severe impairment of the GA response. Quantitative real-time PCR (qRT-PCR) measurements of GA response and biosynthesis genes expression revealed a parallel 2.5 to 3-fold reduction in the transcript levels of all three *GID1* GA-receptors in the *er/erl1/erl2* mutant compared to WT (Figure 1G). The GA-receptor genes *GID1A/B* have been reported to be direct ChIP targets of RGA, a major DELLA repressor of GA-signaling, which stimulates *GID1* transcription (Zentella et al., 2007). The *er/erl1/erl2* triple mutant also displayed altered expression of GA biosynthesis genes compared to the WT: a 4-fold reduction of mRNA levels of *KAO1* (*ent*-kaurenoic acid oxidase) and *KO* (*ent*-kaurene oxidase), a 2.5-fold increase of mRNA levels of GA-repressed *GA3ox1* and *GA3ox2* (GIBBERELLIN 3 BETA-HYDROXYLASE 1 and 2), a 2-fold inhibition and 5-fold up-regulation, respectively, of mRNA levels corresponding to the *GA20ox1* and *GA20ox3* genes (Figure 1H). This indicated that the ERf proteins not only influence the expression of GA receptors, but also genes associated with GA biosynthesis. We subsequently found a substantial decrease of bioactive GA_4_ as well as GA_12,_ and GA_24_ intermediates in *er/erl1/erl2* mutant (Figure 1I, Supplementary Figure 3). Counterintuitively, the Western blotting using a specific antibody (Willige et al., 2007) detected reduced levels of RGA in the *er/erl1/erl2* mutant plants (Figure 1J). Our results indicate that the parallel inactivation of all ERf proteins results in co-ordinate deregulation of GA biosynthesis and response pathways in Arabidopsis.

### ERECTA (ER) Protein Undergoes Endocytosis and Migrates to the Nucleus

In analogy to some human membrane receptors internalizing to endosomes and migrating to the nucleus (*i.e.,* Giri et al., 2005), the ERL2 member of the ERf undergoes endocytosis (Ho et al., 2016). We next examined, in detail, the cellular localization of ER by creating C-terminal GFP fusions with ER (Figure 2A) after verifying genetic complementation of the *er-105* mutation by a 35S::ER-GFP construct (Supplemental Figure 4).

**Figure 2.**
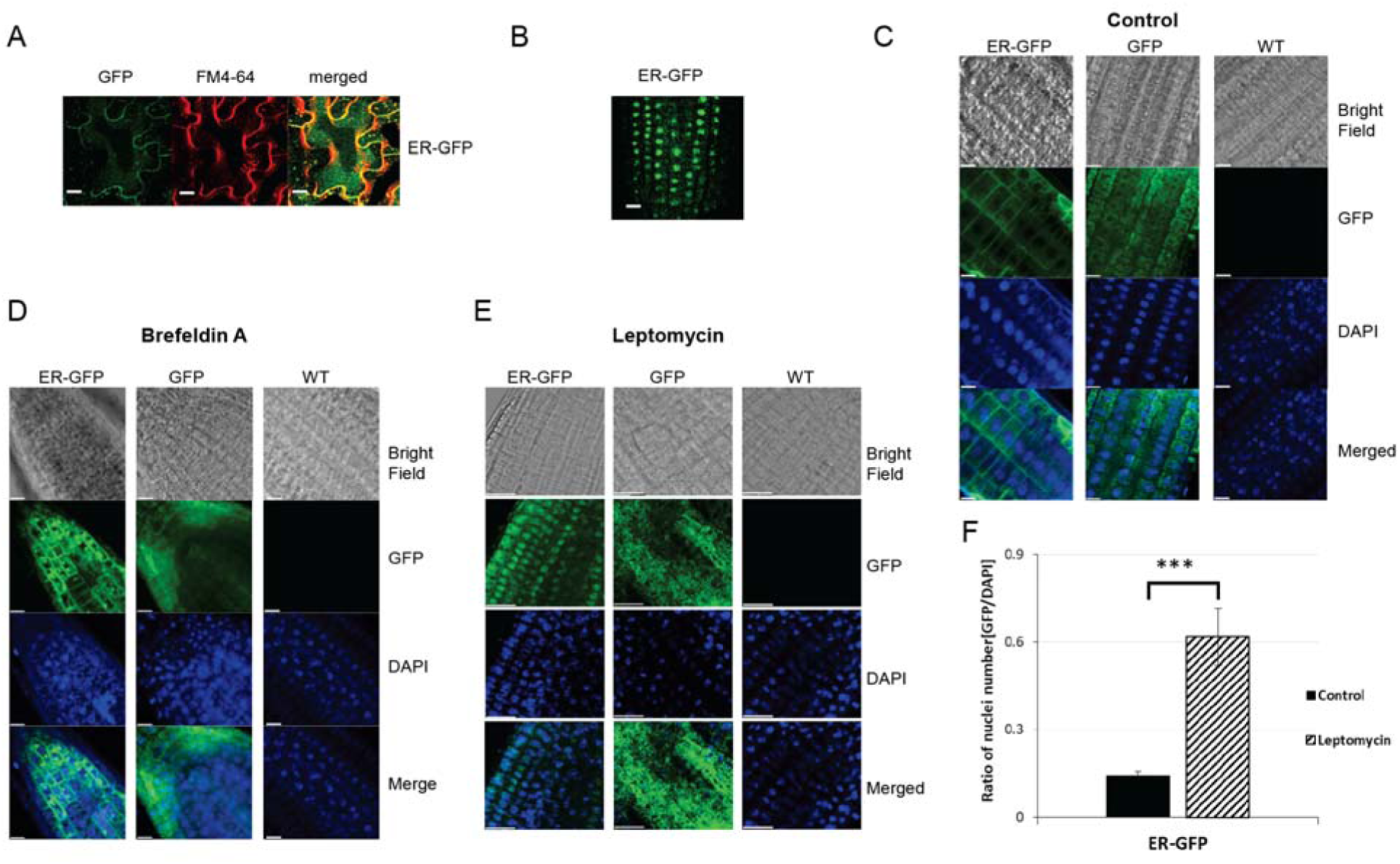
Subcellular localization of ERECTA protein (See also Figures S4 and S5). A, ERECTA is localized in plasma-membrane and endosomes in epidermal cells of 7-days old seedlings. ER-GFP, or free GFP visualized using GFP channel. FM4-64 specifically stains plasma-membranes. Scale bar=10µm. B, Root-tip images of approximately two-week-old (14-17 days) ER-GFP seedlings showing nuclear localization of ERECTA protein at considerable frequency. C, Root-tip images of 12-day-old ER-GFP seedlings serving as the control for D and E. D, Brefeldin A treatment enhanced the localization of ERECTA protein in Brefeldin A (BFA) bodies. Roots of 12-day-old Arabidopsis seedlings. E, Leptomycin B treatment enhanced the nuclear localization of ERECTA Free GFP was used as a control in C, D, and E, cell nuclei were stained with DAPI, scale bar= 50µm. F, Letomycin B enhances nuclear presence of ER protein. The GFP/DAPI ratio calculated per area for roots of plants expressing ER-GFP protein.

As observed earlier (Shpak et al., 2005; Uchida et al., 2012a), a pool of the ER-GFP protein was detected in association with plasma-membranes of the leaf epidermis. In addition, a weak localization signal was detected in internal structures, which could represent endosomes (Figure 2A). In guard cell pairs, ER-GFP protein was also detected in circles around the positions of nuclei, which were visualized by propidium iodide staining (Supplemental Figure 5A and Supplemental Movie 1). We also observed with considerable frequency ER protein in the nuclei of roots of 14- to 17-day-old Arabidopsis plants (Figure 2B), however ER was mainly located in the plasma membrane and endosome-like structures (Figure 2C).

To verify that the ER-GFP protein indeed undergoes endocytosis, we examined its localization in Arabidopsis seedlings treated with 25µM Brefeldin A (BFA), a compound preventing Golgi-mediated vesicular transport of membrane proteins to the plasma membrane (Miller et al., 1992). We observed accumulation of the ER-GFP protein in BFA bodies within 30-40 min after BFA treatment leading to its accumulation at the nuclei periphery 120 min after BFA application (Figure 2D, Supplemental Figure 5B). The 4h long 200 nM leptomycin B treatment (a compound blocking nuclear export by EXPORTINS (Haasen et al., 2002)) resulted in the ER accumulation in the cell nuclei (Figure 2E, F). ER thus appeared to behave similarly to certain human plasma-membrane receptors in migrating into the nucleus (Hung et al., 2008; Chen and Hung, 2015).

We noted that ERL1 and ERL2 proteins carry a monopartite nuclear localization signal (NLS) sequence in their kinase domains, while the ERL1 kinase domain carries an additional bipartite NLS identified using cNLS Mapper (Kosugi et al., 2009b). The NLS signal in the ER protein was not recognized, however, all ERf proteins show evolutionally conserved amino acid sequences in this region (Supplemental Figure 6A). Using the NetNES1.1 (la Cour et al., 2004) server, we predicted the existence of specific for AtXPO1/AtCRM1 exportin (Haasen et al., 2002) leucine-rich nuclear export signals (NES) in all ERf proteins (Supplemental Figure 6A). The subsequent Western-blotting analysis of nuclear extracts (Supplemental Figure 6B) confirmed the nuclear presence of ER. In addition to the expected full-length forms (140 kDa), we also detected shorter (∼75 kDa) versions of the ERECTA protein with the C-terminal GFP tag and smaller products of degradation, including free GFP, suggesting an analogy to the human Epidermal Growth Factor Receptor (EGFR), (Chen and Hung, 2015). The detection of N-terminally truncated forms of the ERECTA protein carrying a kinase domain resembled the recently reported fate of the XA21 LRR-RLK immune receptor in rice (Park and Ronald, 2012), where its C-terminal kinase domain enters the nucleus to interact with the OsWRKY62 transcriptional regulator.

To assess whether the ERECTA kinase domain (KDER) is imported into the nucleus, we fused the C-terminal part of ERECTA, harboring the KDER, to a YFP-HA tag (Supplemental Figure 6C) and expressed this construct in the *er-105* mutant. The presence of KDER-YFPHA was detected exclusively in cell nuclei (Figure 3A). Furthermore, KDER-YFPHA expression partially restored the *er-105* rosette and cauline leaf phenotype to WT values (Supplemental Figure 7A, B). Still, it failed to genetically complement the defect of stem elongation (Supplemental Figure 7C). Partial genetic complementation of the *er-105* mutation and nuclear localization of KDER prove that the KDER has a receptor-domain independent signaling function. Interestingly, by contrast to the full length and kinase domain of ERECTA protein (Figure 3A), truncated ERΔkinase form of ERECTA protein fused to YFP-HA (ΔKDER-YFP-HA) did not enter into the nucleus proving the presence of functional NES and NLS sequences in the C-terminal part of ERECTA protein.

**Figure 3.**
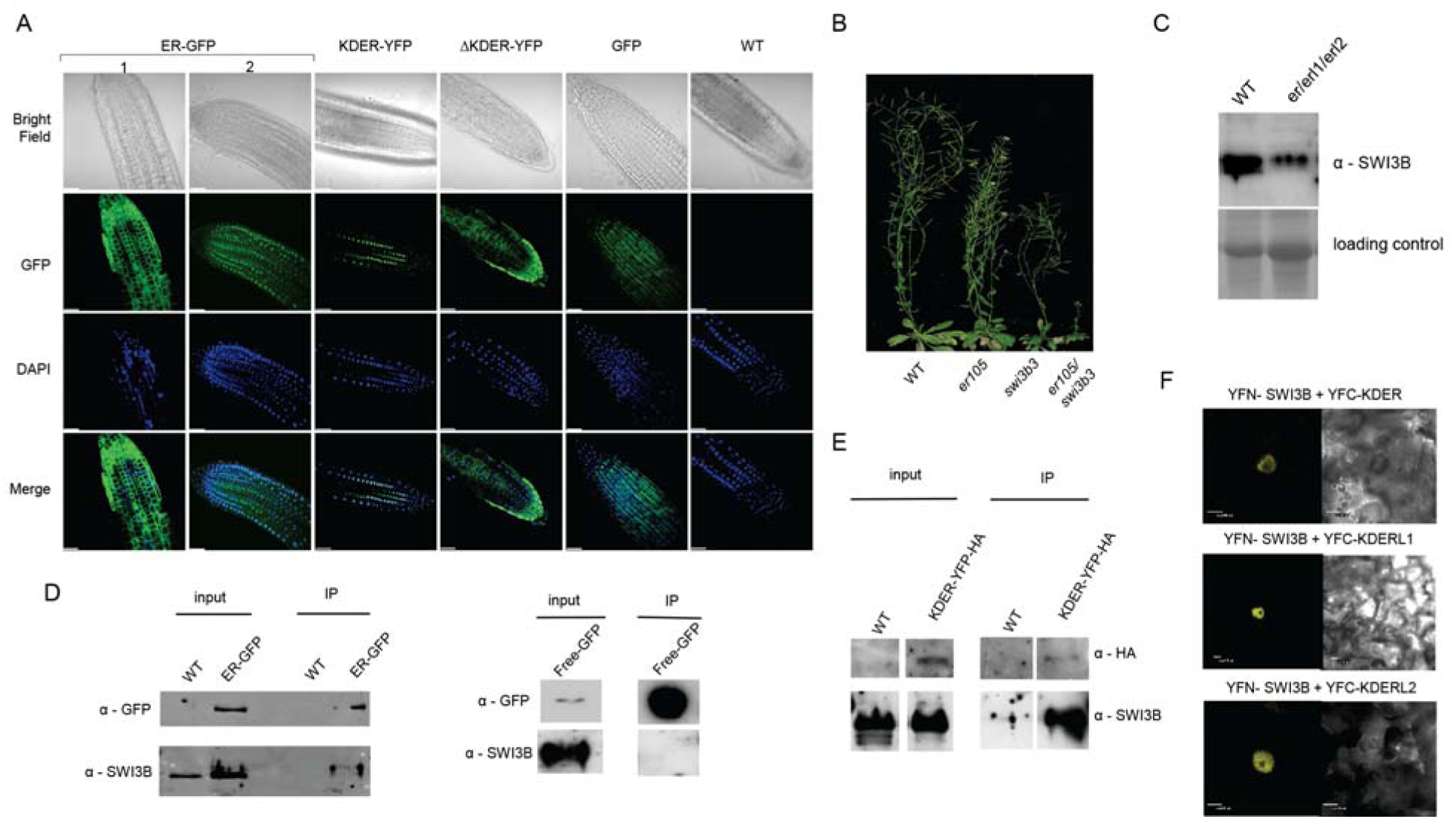
Nuclear function of ERf proteins (See also Figures S6, S7, S8, S9, and S10). A, Root-tip images of approximately two-week-old (14-17 days) plants expressing ER-GFP, KDER-YFP-HA (the kinase domain of the ER protein), ERΔK-YFPHA (truncated ER protein lacking the kinase domain) proteins indicating that the kinase domain is necessary for the nuclear localization of ER protein. WT and GFP expressing plants-negative controls. Panel 2 in the ER-GFP indicates nuclear localization of ER protein appearing at considerable frequency. Cell nuclei were stained with DAPI. Scale bar=25 µm. B, The *er-105/swi3b3* double mutant shows more retarded growth than either *er-105* or *swi3b3* plants. Scale bar= 1cm. C, The *er/erl1/erl2* triple mutant exhibits reduced SWI3B protein level. D, ER-GFP or free GFP (negative control) pull-down from the nucleus and anti-SWI3B western blotting indicate a specific ER-SWI3B interaction. E, Immunoprecipitation of KDER-YFP-HA from the nucleus indicated that the kinase domain of ER interacts with SWI3B. F, ER, ERL1, and ERL2 kinase domains interact with SWI3B in the nucleus. Bimolecular Fluorescence Complementation assay (BiFC) in epidermis of tobacco leaves. Scale bar = 10µm.

Upon nuclei fractionation (Sarnowski et al., 2002), the ERECTA protein was detected in the nuclear membrane, soluble nuclear-protein fraction, and chromatin, with its major presence within the nuclear matrix. The nuclear fractions contained both full-length and N-terminally truncated ER forms (Supplemental Figure 8) suggest that ER could be involved in either transcriptional regulation or other nuclear functions.

### ERf Proteins Physically Interact with the SWI3B Core Subunit of SWI/SNF CRC

The weak *swi3b-3* allele (Sáez et al., 2008) carrying, in the *er-105* background, a point mutation in the *SWI3B* gene encoding a core subunit of the SWI/SNF chromatin remodeling complex (CRC) exhibit severe dwarfism, altered leaf shape, delayed flowering and reduced fertility (Figure 3B). We, therefore, introgressed the *swi3b-3* mutation into WT and found that the phenotypic alterations related to *swi3b-3* were much weaker (slight reduction of growth rate, leading to decreased plant height, Figure 3B) than the phenotypic traits exhibited by the *er-105*/*swi3b-3* as well as single *er-105* mutation. The severe phenotypic alterations exhibited by the *er-105*/*swi3b-3* plants indicated the likely existence of a strong genetic interaction between the ERECTA signaling pathway and SWI3B-containing SWI/SNF CRCs. This observation is in line with *i)* the direct binding of 15 out of 27 potential ERf target genes related to the GA signaling pathway (Supplemental Table 1) by SWI/SNF CRCs (Sacharowski et al., 2015; Archacki et al., 2016; Li et al., 2016); *ii)* the unexpected broad transcriptional changes and severe effects on Arabidopsis development and hormonal signaling pathways observed in the *er/erl1/erl2* mutant, and *iii)* the well-recognized function of SWI/SNF CRC in hormonal crosstalk including GA signaling (Sarnowska et al., 2013; Sarnowska et al., 2016).

We next assessed the level of the SWI3B protein in *er/erl1/erl2* plants. We found a significant decrease in the SWI3B protein abundance (Figure 3C), further suggesting that the ERf signaling pathway may influence the proper function of SWI3B-containing SWI/SNF CRCs. Additionally, the SWI3B was found to bind ER-GFP but not free GFP (Figure 3D). Similarly, co-immunoprecipitation indicated that SWI3B interacts with the kinase domain of ER (Figure 3E).

Next, we performed BiFC assays (Hu et al., 2002) in epidermal cells of *Nicotiana benthamiana* and confirmed the SWI3B and ER kinase domain interaction. The YFC-RFP served as a control unrelated protein with broad intracellular localization (Figure 3F, Supplemental Figure 9). We also detected the interaction of SWI3B with the kinase domain of the ERL1 or ERL2 (Figure 3F, Supplemental Figure 9), indicating the existence of direct interdependences between the ERf signaling pathway and SWI/SNF–dependent chromatin remodeling.

Moreover, we found similar interactions in the nuclei of human cells for HER2 (Epidermal Growth Factor Receptor-family member), a membrane receptor acting in a non-canonical signaling mode including translocation to the nucleus (Lee et al., 2015a), and BAF155 a SWI3-type subunit of human SWI/SNF CRCs (Supplemental Figure 10). Thus, our data indicate that the phenomenon observed for ERf and SWI3B is not limited to Arabidopsis but rather may be a general feature of SWI/SNF CRCs and membrane receptors.

### *ERECTA* and *SWI3B* Interact Genetically and *er/elr1/erl2* Plants Exhibit Alteration in Chromatin Status

The *er-105/swi3b-3* double mutant exhibited more severe phenotypic traits than both single *er-105* and *swi3b-3* mutant lines (Figure 4A), supporting the observed physical interdependences between ER and SWI3B.

**Figure 4.**
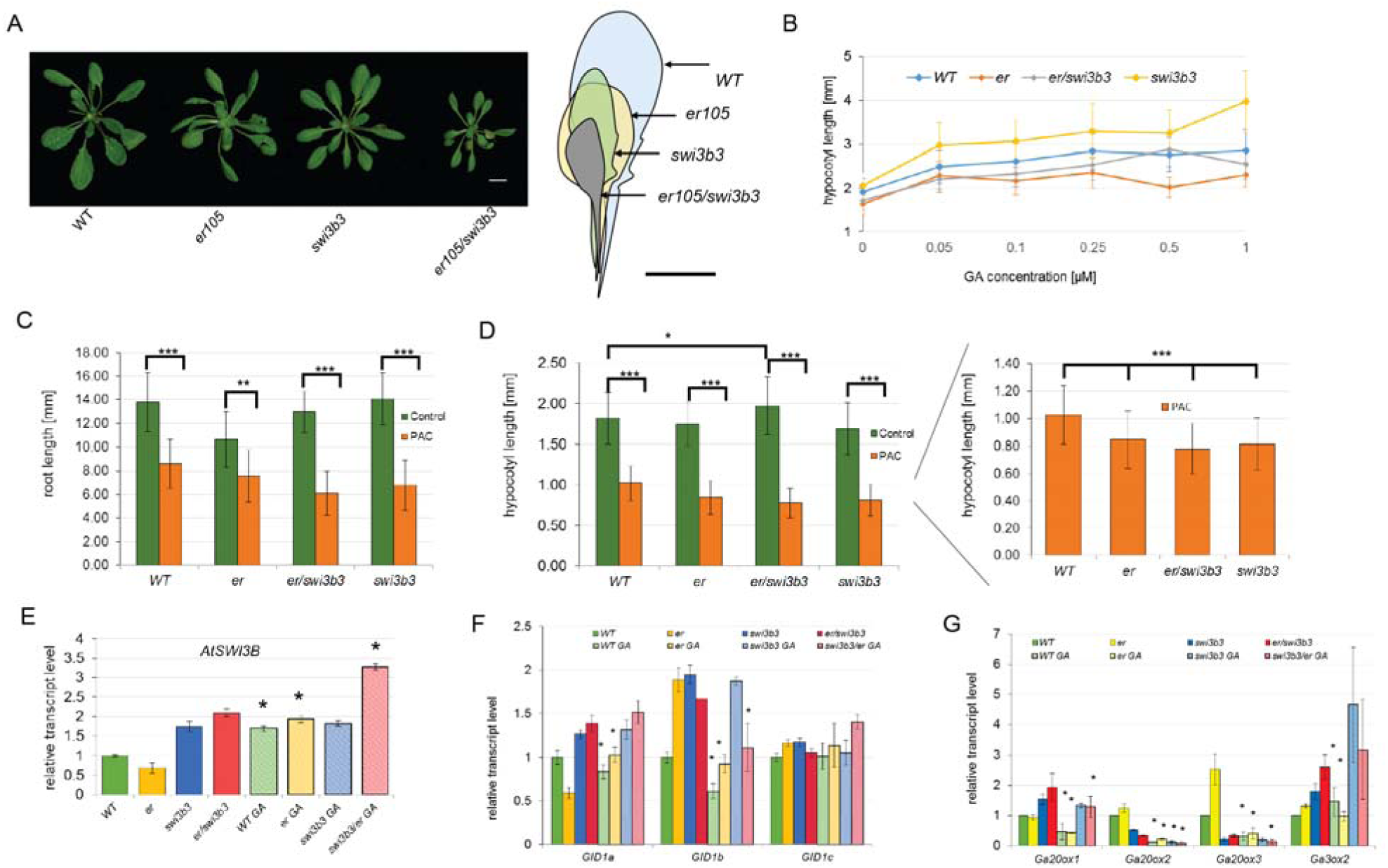
ER and *SWI3B* interact genetically and affect both GA biosynthesis and response pathways (See also Figures S11 and S12). A, The *er-105/swi3b-3* double mutant exhibits more retarded growth than the *er-105* and *swi3b-3* (three-weeks old plants). Graphical alignment of corresponding leaves. Scale bar= 1 cm. B, The hypersensitivity of 1-week-old *swi3b-3* hypocotyl to GA treatment is abolished by introducing *er-105*. C, Roots of all tested 1-week-old genotypes similarly respond to PAC treatment (error bars-SD, *P < 0.01,** P < 0.001, ***P<0.0001, Student’s *t*-test). D, Hypocotyls of all tested 1-week-old genotypes similarly respond to PAC treatment, right panel-hypocotyl length comparison for PAC treated plants only (error bars-SD, *P < 0.01,** P < 0.001, ***P<0.0001 Student’s *t*-test). E, *swi3b-3* weak, point mutant line and *er-105/swi3b-3* exhibit elevated *SWI3B* transcript level, the *SWI3B* expression is elevated after supplementation with bioactive GA_4+7_ in all genotypes except *swi3b-3* (error bars-SD, P < 0.05, Student’s *t*-test). F, The examination of *GID1* genes indicated that almost all examined lines responded to GA treatment, but the *swi3b-3* line was insensitive for GA-induced transcriptional changes (error bars-SD, P < 0.05, Student’s *t*-test). G, The examination of GA biosynthesis genes indicated that almost all examined lines responded to GA treatment, but the *swi3b-3* line was insensitive for GA-induced transcriptional changes except *GA20ox2* expression (error bars-SD, P < 0.05, Student’s *t*-test).

The treatment with bioactive GA_4+7_ gibberellins indicated hypersensitivity of *swi3b-3* to GA demonstrated by hypocotyl length, while the response of *er-105* was reduced. By contrast, the GA hypersensitivity of *swi3b-3* was abolished by introduced *er-105* mutation (Figure 4B). All tested genotypes were responding to PAC treatment in a similar way (Figure 4C, D). The higher expression of *SWI3B* was visible in the case of the *swi3b-3* mutant, which was even more pronounced in *er-105/swi3b-3*. The expression of *SWI3B* was elevated after supplementation with bioactive GA_4+7_ in all genotypes, except for the *swi3b-3* line (Figure 4E). The examination of *GID1* and GA biosynthesis genes indicated that almost all examined lines responded to GA treatment while the *swi3b-3* line was insensitive for induced by GA transcriptional changes except for *GA20ox2* expression (Figure 4F, G). Collectively, our results further indicate that both ERf signaling and SWI3B-containing CRCs play together an important role in the fine-tuning of GA signaling in Arabidopsis.

To verify the biological effect of observed interactions between ERf signaling and SWI3B-SWI/SNF, we analyzed the chromatin status, nuclei shape and chromocenters number in the *er/erl1/erl2* mutant plants. We found that *er/erl1/erl2* plants exhibit increased chromocenter number and altered spindle-like nuclei shape (Supplemental Figure 11A, B). We furthermore screened the effect of inactivation of ERfs on genome-wide nucleosome positioning in chromatin using *micrococcal nuclease* protection assays followed by deep sequencing (MNase-seq) and confirmatory MNase-qPCR in WT and *er/erl1/erl2* plants.

We found that inactivation of ERf proteins has a broad influence on the global nucleosomal chromatin structure in Arabidopsis-*erf* exhibited 41519 nucleosome occupancy changes, 13924 “fuzziness” changes and 4055 nucleosome position changes (Supplemental Figure 12A) and alterations in the presumable regulatory regions upstream of the transcription start site (TSSs) (Supplemental Figure 12B, Supplemental Dataset 1, Sub-Table 9-14).

Among genes with down-regulated expression and altered nucleosome positioning in the *er/erl1/erl2* mutant were 14 GA-related genes (*ATBETAFRUCT4, XERICO, PRE1, MYBR1, MYB24, MIF1, HAI2, ZPF6, GA20ox1, CGA1, XTH24, GID1b, RGL1,* and *GIS3*) Interestingly, seven of them (*ATBETAFRUCT4, PRE1, MYBR1, MIF1, XTH24, GID1b,* and *RGL1*) were already observed to be directly targeted by the BRM ATPase of the SWI/SNF CRC (Archacki et al., 2016; Li et al., 2016).

An Integrated Gene Browser (IGB) view of *PRE1, GID1a,b* promoter regions indicated (Supplemental Figure 12D) various nucleosome alterations on promoter regions of these genes in the *er/erl/erl2* mutant pointing out impaired chromatin remodeling in the absence of functional ERf proteins. The selected changes were confirmed by MNase-qPCR (Supplemental Figure 12E).

### The Inactivation of ERf proteins Affects SWI3B Protein Phosphorylation

We tested the ability of KDER to phosphorylate SWI3B protein. We overexpressed, purified, and subsequently used MBP-His6-KDER and His6-SWI3B (Figure 5A, Supplemental Figure 13A) for non-radioactive *in vitro* kinase assay. The existence of a strong band corresponding to phosphorylated SWI3B protein and a weaker band of autophosphorylated KDER was indicated (Figure 5B, Supplemental Figure 13B, C). The confirmatory mass-spectrometry analysis resulted in the identification of the active phosphorylation sites at KDER and in the SWI3B protein (Supplemental Figure 13D, E). Interestingly three of four KDER- dependent phosphorylation sites were located in SWI3B in SWIRM and SANT domains (Supplemental Figure 13E, F), providing a valuable hint that the ERf family proteins may be responsible for the SWI3B phosphorylation.

**Figure 5.**
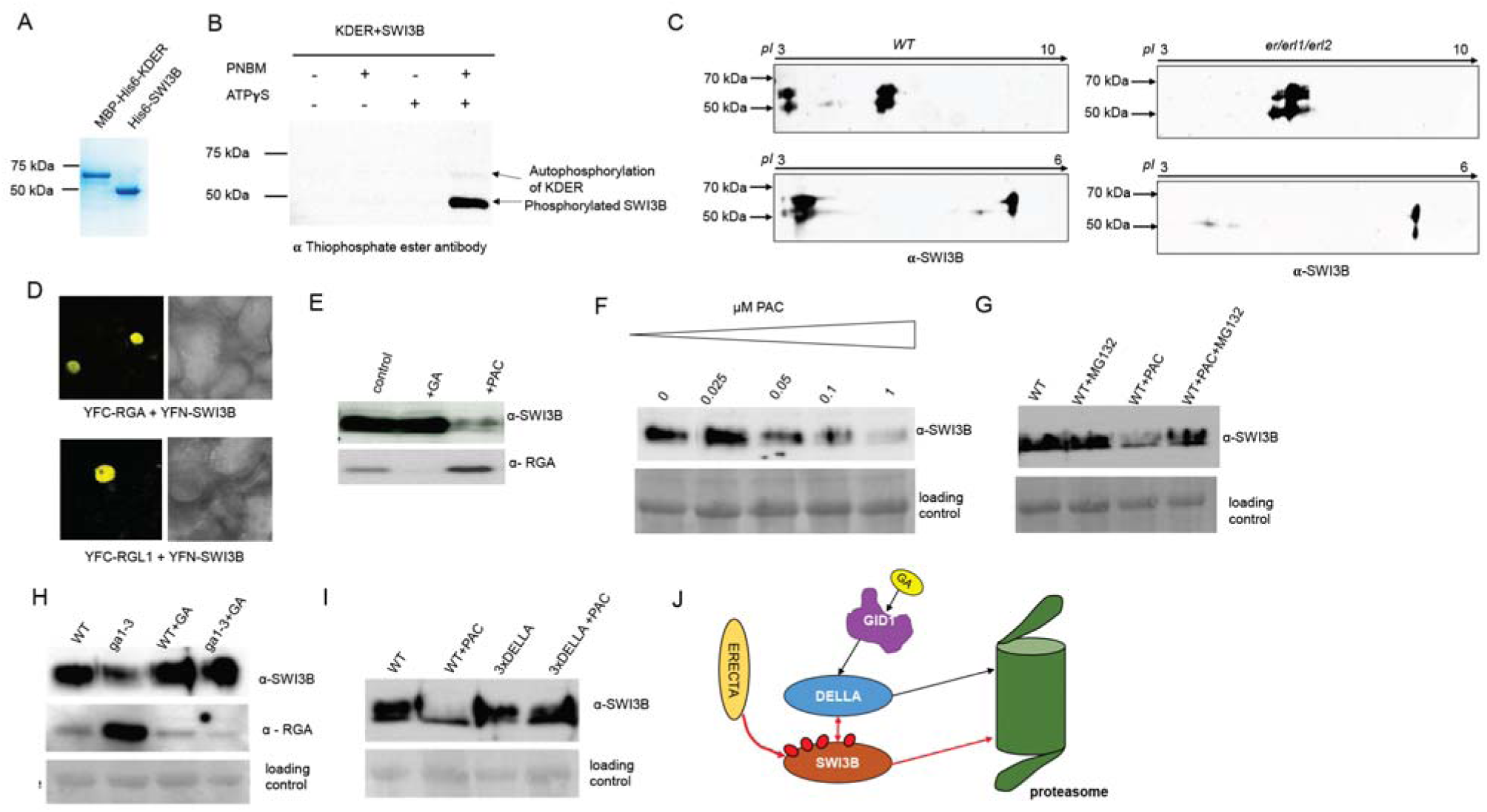
ERf proteins are responsible for the phosphorylation of SWI3B protein, while DELLA proteins control SWI3B protein abundance (See also Figures S13, S14, and S15). A, Coomassie staining of MBP-His6-KDER and His6-SWI3B proteins purified from bacteria. B, Western blot with anti-Thiophosphate ester antibody (ab92570; Abcam) showing *in vitro* SWI3B phosphorylation by KDER. C, 2D Western blot assay with anti SWI3B antibody indicating *in vivo* phosphorylation alteration of SWI3B protein in *er/erl1/erl2* mutant. D, SWI3B and RGA and RGL1 proteins in the nuclei of living cells. Bimolecular Fluorescence Complementation assay (BiFC) in epidermis of tobacco leaves. Scale bar = 10µm. E, The amounts of SWI3B and RGA proteins in plants are oppositely regulated by PAC treatment. F, The disappearance of SWI3B protein is PAC-dose dependent. G, The PAC-dependent degradation of SWI3B is abolished by the MG132 treatment, a known proteasome inhibitor. H, The *ga1-3* mutant constitutively accumulating DELLA proteins exhibits the decreased level of SWI3B, which is restored to WT levels upon GA treatment. I, The triple DELLA mutant exhibits a WT-like level of SWI3B protein, and the PAC treatment does not influence SWI3B level in this background. J, Schematic model highlighting ERf and DELLA impact on the SWI3B protein.

To verify this possibility the *in vivo* phosphorylation analysis using 2D-IEF-PAGE combined with the Western blot was performed. The alteration in the SWI3B proteins isoelectric point (*pI*) in *er/erl1/erl2* mutant was observed. The bands corresponding to phosphorylated SWI3B form were nearly absent in *er/erl1/erl2* plants, indicating a severe defect in SWI3B phosphorylation (Figure 5C). This data strongly supports the regulatory function of the ERf proteins on the SWI3B subunit of SWI/SNF CRC.

### Accumulation of DELLA Proteins Correlates with Increased Proteasomal Degradation of SWI3B

Our previous study demonstrated that SWI3C, a partner of SWI3B, physically interacts with DELLA proteins (Sarnowska et al., 2013). We also found that the Arabidopsis lines with impaired SWI/SNF CRCs-*brm* and *swi3c* exhibit decreased level of bioactive GA_4_ gibberellins level (Sarnowska et al., 2013; Archacki et al., 2013), but they do not accumulate RGA DELLA protein similarly as in case of *er/erl1/erl2* plants (Supplemental Figure 14 and Figure 1J). To address this unusual phenomenon, we used a BiFC assay to analyze the interaction between DELLA and SWI3B. The interaction between either RGA or RGL1 protein and SWI3B was found (Figure 5D, Supplemental Figure 15). No YFP signal was detected in control cells.

To understand the functional consequences of the detected interactions between SWI3B and DELLA proteins, we analyzed the amounts of SWI3B and RGA proteins in GA or PAC-treated plants (Figure 5E). Surprisingly, we observed the PAC-dose-dependent disappearance of SWI3B protein (Figure 5F). To check if the degradation of SWI3B under these conditions depended on the proteasome, we tested the effect of MG-132 on the SWI3B level. MG-132 treatment caused increasing SWI3B abundance in PAC treated plants (Figure 5G), suggesting that the degradation of SWI3B observed in parallel to accumulation of DELLAs occurs *via* the proteasome. We also observed increased degradation of SWI3B in the *ga1-3* mutant in which DELLA proteins are constitutively accumulated (Figure 5H), but we did not observe enhanced SWI3B degradation in PAC-treated 3xDELLA (Archacki et al., 2013) collectively suggesting that binding of SWI3B by DELLA proteins may be a primary cause of its proteasomal degradation (Figure 5I, J). Thus, the accumulation of DELLA proteins should lead to the same consequences as the elimination of SWI3B protein or SWI3B-containing SWI/SNF CRCs. This conclusion is strongly supported by the lack of RGA protein accumulation in GA deficient *brm* and *swi3c* lines with inactivated other subunits of SWI/SNF CRC (Archacki et al., 2013; Sarnowska et al., 2013). Therefore, it could be indeed expected that GA and SWI3B-deficient *er/erl1/erl2* mutant will also not accumulate RGA protein because, in the case of SWI/SNF CRC impairment, the DELLA accumulation seems to be irrelevant. Collectively, our results provide new insight into the functioning of the ERECTA family proteins and DELLA proteins and their mutual impact on the SWI3B-containing SWI/SNF CRCs.

### Binding of ERECTA and SWI3B to Promoter Regions of the *GID1* Genes

The *er/erl1/erl2* mutant displays an impaired response to exogenous GA treatment and a consistently decreased expression of all three *GID1* genes. ERf proteins interact with the SWI3B, inactivation of *ERf* proteins results in nucleosomal chromatin structure alterations and decreased abundance of SWI3B and its phosphorylated form in Arabidopsis, and there is an intriguing interdependence between the control of the SWI3B level and DELLA protein accumulation, therefore we examined if ER and the SWI3B participate in transcriptional control of the *GID1* genes previously reported as targets for DELLA (Rosa et al., 2015). The ChIP analysis on *GID1* promoters was performed using nuclei purified from 3-week-old seedlings expressing the ER-GFP and the SWI3B-HIS-STREP-HA proteins.

The binding of ER-GFP was detected around -130bp upstream of the TSS in the *GID1a* promoter region while SWI3B bound around the TSS of the *GID1a* promoter (Figure 6A). The ER protein was targeted to two regions around -150bp and -350bp from the TSS in the *GID1b* promoter, SWI3B was localized only -350bp upstream TSS (Figure 6B). ER and SWI3B were similarly cross-linked to the -100bp region of the *GID1c* promoter, but SWI3B was also mapped further upstream to -800bp (Figure 6C).

**Figure 6.**
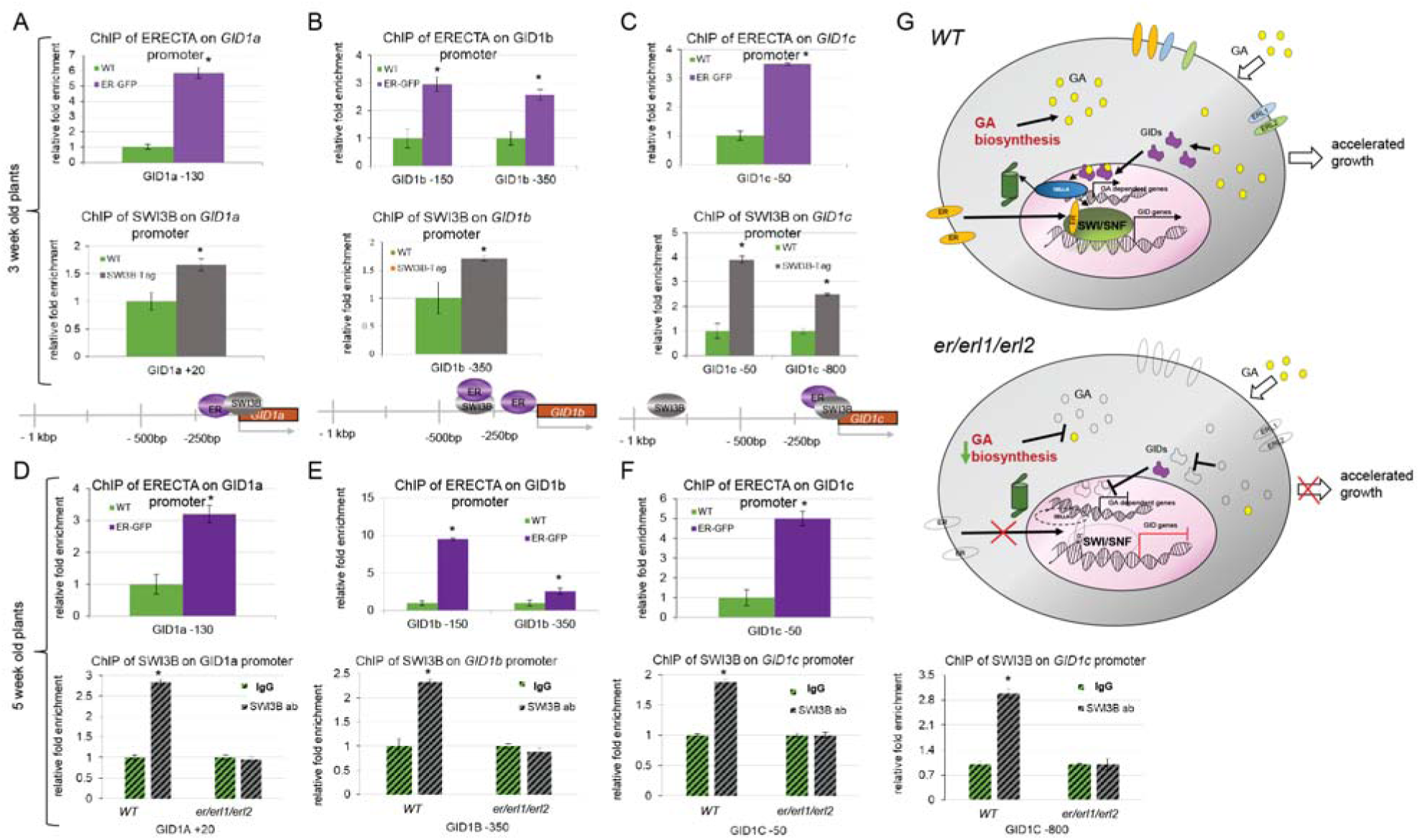
ERf proteins enter the nucleus where ERECTA protein binds the *GID1* promoters similarly to the SWI3B subunit of SWI/SNF CRC (See also Figures S16 and S17). A, ERECTA protein binds to promoter regions of the *GID1a* gene in a region targeted by the SWI/SNF complex in three-week-old plants. (error bars refer to SD, P < 0.05, Student’s *t*-test, three biological and technical replicates were performed). B, ERECTA and SWI3B core subunit of SWI/SNF CRCs target promoter regions of *GID1b* gene in three-weeks old plants (error bars refer to SD, P < 0.05, Student’s *t*-test, three biological and technical replicates were performed). C, SWI3B binds to the promoter region of the *GID1c* gene in two different regions. One of them is targeted by ERECTA protein in three-week-old plants. D, ERECTA protein binds to promoter regions of the *GID1a* gene in a region targeted by the SWI/SNF complex in five-week-old plants. (error bars refer to SD, P < 0.05, Student’s *t*-test, three biological and technical replicates were performed). E, ERECTA targets promoter region of *GID1b* gene in five-week-old plants (error bars refer to SD, P < 0.05, Student’s *t*-test, three biological and technical replicates were performed). F, ERECTA binds to the promoter region of the *GID1c* gene in three-week-old plants. The bottom panel in D-F: the binding of native SWI3B protein to its target sites in *GID1a-c* promoter regions is abolished in 5-week-old *er/erl1/erl2* triple mutant plants. G, A model describing the non-canonical nuclear function of ERf proteins in the GA signaling pathway.

Inspection of ER and SWI3B binding to the *GID1a-c* promoter regions in 5-week-old WT, ER-GFP, and the *er/erl1/erl2* mutant demonstrated that ER binds the same *GID1* promoter regions as in the case of 3-week-old plants (Figure 6 D-F). The SWI3B binding to the promoter of *GID1* genes was abolished by the inactivation of ERf proteins in *er/erl1/erl2* plants. The inactivation of *ER, ERL1,* or *ERL2* did not affect the binding of SWI3B to *GID1a-c* promoter regions in single *er105, erl1,* and *erl2* mutant lines (Supplemental Figure 16). Our study provides evidence that the three ERf proteins have redundant functions regarding proper SWI3B recruitment since only the simultaneous absence of all ERfs proteins abolished SWI3B binding to the *GID1a-c* promoters.

Of note, we found a similar binding of HER2 (EGFR-family) receptor and BAF155 subunit of SWI/SNF CRCs to the promoter regions of human *BRCA1* and *FBP1* genes (Supplemental Figure 17), indicating that the phenomenon observed for ER may be a general mechanism controlling gene expression that is maintained between kingdoms.

## Discussion

Inactivation of ERf LRR-RLK family members results in various defects in Arabidopsis growth and development. While it is well established that ERf proteins play distinct roles in the control of epidermal patterning, stomatal development, meristem size, inflorescence architecture, and hormonal signaling, the exact mechanisms underlying the regulatory functions of ERf proteins in these processes are largely unknown (*e.g.,* Chen and Shpak, 2014; Chen et al., 2013b; Qi et al., 2004; Van Zanten et al., 2010; Kosentka et al., 2019).

Here we show that the inactivation of Arabidopsis ERf proteins has a broad effect on various regulatory processes, including hormonal signaling, and suggest that these sum responses underlie the severe developmental defects exhibited by the *er/erl1/erl2* mutant. We demonstrate that parallel inactivation of all *ERf* proteins results in severe deregulation of the GA signaling pathway as evidenced by the impairment of GA perception and GA biosynthesis. When taken together, these findings, alongside the identification of NLS and NES sequences in ERf proteins and our demonstration of their translocation into the nucleus, suggest a novel, non-canonical function of ERf proteins (Figure 6G).

Our study also reveals an analogy of this system to the previously described XA21 LRR immune receptor in rice (Park and Ronald, 2012) and to the non-canonical signaling mode of the human Epidermal Growth Factor Receptor (EGFR) family (Lee et al., 2015a). Although it should be stressed that these two classes of plant and animal epidermal receptors carry completely unrelated sequences from one another and from the system we describe here (Supplemental Table 3), suggesting that the translocation of the membrane receptors to the nucleus may be a general paradigm maintained between plant and animal kingdoms. In addition to their canonical membrane receptor functions, holoreceptor and truncated forms of EGFRs are imported into nuclei *via* ER-mediated retrograde transport, although some of them lack known NLSs (Chen and Hung, 2015).

In the nucleus, the EGFR receptors can bind to DNA, interact with various transcription factors. Thereby, nuclear forms of EGFRs are implicated in the control of cell proliferation, DNA replication and repair, and transcription (Chen and Hung, 2015), so their functions extend far beyond the regulation of epidermal patterning.

We found here that the Arabidopsis ERECTA LRR-RLK receptor similarly translocates from the plasma membrane into the nucleus. Both intact and N-terminally truncated forms of ERECTA were detectable in the nucleus. A truncated ERECTA carrying only the kinase domain localizes exclusively in the nucleus and partially complements the leaf developmental defects caused by the *er-105* mutation implying a ligand-independent non-canonical signaling function of the ERECTA kinase domain.

Furthermore, our data show that the ERf proteins interact through their kinase domains with the SWI3B, and ER can phosphorylate the SWI3B core subunit of the SWI/SNF CRC. SWI/SNF plays a pivotal role in the hormonal crosstalk regulation in both humans and plants (Sarnowska et al., 2016). Moreover, we show that analogously as ER protein, the HER2 member of the EGFR family directly interacts with the BAF155 subunit of human SWI/SNF and co-localizes with BAF155 on some gene promoters providing evidences that such system is likely maintained between kingdoms.

Parallel inactivation of all ERf proteins results in alterations of genome-wide nucleosomal chromatin structure and altered transcriptional activity of a large number of genes. The binding of SWI3B to its target regions in the *GID1a-c* promoters is retained in *er-105, erl1,* and *erl2* single mutant lines. By contrast, the *er/erl1/erl2* mutant plants exhibited a reduction in phosphorylation SWI3B protein level and abolished proper SWI3B binding to *GID1* promoter regions together with decreased expression of *GID1a-c* genes, indicating a strong and direct effect of the ERf signaling pathway on SWI/SNF-dependent chromatin remodeling. The *er/erl1/erl2* mutant plants are characterized by a decreased level of endogenous gibberellins; however, they do not accumulate DELLA proteins, similar to the SWI/SNF mutants. Thus, we have demonstrated that the *er/erl1/erl2* mutant plants exhibit severe deregulation of the gibberellin signaling pathway and SWI/SNF-dependent chromatin remodeling. This is, in turn, an attractive explanation of the observed insensitivity of *er/erl1/erl2* mutant to the application of exogenous gibberellin. DELLA proteins are involved in the sequestration of various transcription factors and chromatin remodeling complexes (Phokas and Coates, 2021). In this study, we extend the existing knowledge on DELLA functioning by providing evidence for the existence of the DELLA-SWI3B regulatory module and explaining why some GA-deficient mutant lines with impaired SWI/SNF chromatin remodeling complex do not accumulate DELLA proteins (Figure 5G).

Collectively our finding that plant ERf proteins play an important, non-canonical nuclear function, *i.e.,* bind directly to chromatin and control the proper recruitment of SWI/SNF CRCs, which are strongly involved in controlling regulatory processes including hormonal crosstalk (Sarnowska et al., 2016), may be a general paradigm for other classes of plant and mammalian membrane receptor kinases.

## Methods

### Plant Material and Growth Conditions

The *Arabidopsis thaliana* ecotype Columbia was used as wild type (WT) in all experiments. The following Arabidopsis mutants were used for analysis: *er-105*, *er-105/swi3b-3* (Sáez et al., 2008), *er/erl1/erl2* plants (Shpak et al., 2004) and *erf* lines in various combinations (Torii et al., 1996), the 35S::GFP Arabidopsis line has been obtained from NASC (N67775). Seeds were sown on soil or plated on ½ Murashige and Skoog medium (Sigma-Aldrich) containing 0.5% sucrose and 0.8% agar. Plants were grown under long day (LD) condition (12h Day/12h Night or 16h Day/8h Night). For GA response tests, plants were sprayed twice a week with 100 µM GA_4+7_ or water (control) for a fast response the 2h of GA_4+7_ treatment was performed.

### Construction of Transgenic Lines

Genomic sequences of *ERECTA*, cDNAs of *ERECTA* kinase domain and truncated ERECTA lacking kinase domain (ERΔK) were cloned into binary vector p35S::GW::GFP (F. Turck, Max-Planck-Institut für Züchtungsforschung, in the case of ERECTA), and into pEarley Gate 101 (in the case of the ER kinase domain and ΔKDER; (Earley et al., 2006). Plants were transformed using *Agrobacterium tumefaciens* GV3101 (pMP90) by floral-dip method (Davis et al., 2009). The STOP codon of the *SWI3B* genomic sequence was replaced with HIS-STREP-HA using the recombineering method (Bitrián et al., 2011), moved into pCB1 vector (Heidstra et al., 2004), and transformed into *swi3b-2* Arabidopsis mutant line.

### RNA Extraction and qRT-PCR Analysis

Total RNA was isolated from adult (5-week-old) plants using an RNeasy plant kit (Qiagen), treated with a TURBO DNA-free kit (Ambion). Total RNA (2.5µg) was reverse transcribed using a first-strand cDNA synthesis kit (Roche). qRT-PCR assays were performed with SYBR Green Master mix (Bio-Rad) and specific primers for PCR amplification. Housekeeping genes *PP2A* and *UBQ5* (AT1G13320 and AT3G62250, respectively) were used as controls. The relative transcript level of each gene was determined by the 2^-ΔΔCt^ method (Schmittgen and Livak, 2008). Each experiment was performed using at least three independent biological replicates. qRT-PCR primers are listed in Supplemental Dataset 3.

### Transcript Profiling and Gene Ontology Analysis

RNA was isolated from adult (5-week-old) WT and *er/erl1/erl2* plants using a Plant RNeasy kit (Qiagen) according to the manufacturer’s protocol. Transcriptomes were analyzed using 150ng of total RNA as starting material. Targets were prepared with a cDNA synthesis kit followed by biotin labeling with the IVT labeling kit (GeneChip 39IVT Express; Affymetrix) and hybridized to the ATH1 gene chip for 16h as recommended by the supplier. The raw data were analyzed using GenespringGX according to the manual (guided workflow). GO-TermFinder was used for GO analyses of selected groups of genes (Boyle et al., 2004).

### Nuclear Fractionation

Nuclei were isolated from 2g of leaves of 3-weeks old Arabidopsis seedlings according to the method previously described by Gaudino and Pikaard (1997). Subsequent nuclear fractionation was performed using the high-salt method, with modifications (Sarnowski et al., 2002).

### Protein Interaction Study, Confocal Imaging, Subcellular Localization, Brefeldin A and Leptomycin B Treatment, DAPI Staining

Protein interaction was analyzed by performing the immunoprecipitation of ER-GFP or KDER-YFP-HA from nuclei from 4 g of Arabidopsis plants (Saleh et al., 2008). The nuclear extracts were incubated with 25 µL of GFP Magnetic Trap beads (Chromotek) according to manufactures instructions. The presence of SWI3B protein was determined by western blot analysis using anti-SWI3B antibody (Sarnowski et al., 2002).

The interaction between human proteins was analyzed by immunoprecipitation of HER2 and BAF155 from viscolase treated nuclear extracts prepared, according to Jancewicz et al. 2021. The presence of HER2 and BAF155 was determined by Western blot analysis using anti HER2 (CST, 12760) and anti BAF155 (CST, 11956) antibodies.

To obtain YFN-ERL1, YFC-ERL1, YFC-ERL2, and YFC-KDER fusions for BiFC (Hu et al., 2002) analysis, cDNAs encoding ERL1 and ERL2 proteins and ERECTA, ERL1, and ERL2 C-terminal kinase domains were PCR amplified and cloned into the binary vectors pYFN43 or pYFC43 (Belda-Palazón et al., 2012). The *in vivo* interactions between proteins were detected by BiFC using Leica TCS SP2 AOBS, a laser scanning confocal microscope (Leica Microsystems). Tobacco (*Nicotiana benthamiana*) epidermal cells were infiltrated using *Agrobacterium tumefaciens* GV3101 (pMP90) carrying plasmids encoding ERL1, ERL2, or KDER fusions and the p19 helper vector and analyzed by confocal microscopy 3 d later. YFN-RFP and YFC-RFP fusions were used to detect transformed cells in the BiFC assays (Sarnowska et al., 2013); at least five nuclei were analyzed in three separate experiments.

The vesicle trafficking inhibitor BFA (Sigma Aldrich) was used at the 25µM concentration at the following time points 40 min, 90 min, and 120 min. The NES-dependent nuclear export inhibitor Leptomycin B was used at the 200 nM concentration 4h before microscopy observation. Nuclei were stained with 4’,6-diamidino-2-phenylinodole (DAPI) at the 1µg/mL concentration for 30 min. The observation was carried out on the root tip of about two weeks old plants incubated directly before in ½ MS alone or with the addition of proper compound (BFA or Leptomycin B, respectively). Every time 30 min before the end of incubation, DAPI was added.

### Chromatin Immunoprecipitation

ChIP experiments were performed as described previously (Sacharowski et al., 2015) on three or five-week-old WT, ER-GFP, SWI3B-His-Strep-HA, and *er/erl1/erl2* plants cross-linked under vacuum using formaldehyde (final concentration: 1%) and Bis-(sulfosuccinimidyl) glutarate (final concentration: 1mM). For ER-GFP, chromatin immunoprecipitation was performed with GFP-Trap M (Chromotek). For SWI3B ChIP experiments, NiNTA Agarose (Qiagen) or, in the case of anti-SWI3B antibody, the magnetic protein A and G dynabeads (Dynal) were used. ChIP enrichment was determined using qPCR, and relative fold change was calculated using the 2^-ΔΔCt^ method (Schmittgen and Livak, 2008). The TA3 retrotransposon was used as negative control (Pastore et al., 2011). Primers used in ChIP experiments are listed in Supplemental Dataset 3.

Chromatin from the SKBR-3 human cell line was immunoprecipitated according to (Komata et al., 2014 and Jancewicz et al., 2021). Recovered chromatin was incubated O/N at 4°C with the following antibodies: anti BAF155 (CST, 11956), antiHER2 (CST, 12760), and Normal Rabbit IgG (CST, 2729, mock control). Results were calculated based on the 2^-ΔΔCt^ (Schmittgen and Livak, 2008). The relative fold enrichment of the analyzed sample represents the fold change with reference to IgG (mock) sample. A set of primers used for ChIP-qPCR analysis is listed in Supplementary Dataset 3.

### MNase Mapping of Genome-wide Nucleosome Positioning and MNase-qPCR

The nuclear extraction, MNase treatment, subsequent NGS analyses, and confirmatory MNase-qPCR were performed according to (Sacharowski et al., 2015) on 5 week old plant material.

### Gibberellin Analysis

About 200 mg of frozen materials from 5 week old plants were used to extract and purify the GA as described in Plackett et al. (2012) with minor modifications. GA was quantified using MS/MS analysis using 4000 Triple Quad (AB Sciex Germany GmbH, Darmstadt, Germany) in multiple reaction monitoring (MRM) scan with electrospray ionization (ESI) as described in Salem et al., (2016). The mass spectrometry attached to UPLC system (e.g., Waters Acquity UPLC system, Waters, Machester, UK) separation was achieved on a reversed phase C18-column (100 mm × 2.1 mm 1.8 μm).

### *In vitro* Phosphorylation Analysis and *in vivo* Kinase Assay

SWI3B-6xHis was overexpressed and purified, according to Sarnowski et al. (2002). The KDER (pDEST-6xHis-MBP vector) was purified using tandem purification using MBP and Ni-NTA resins. *In vitro* kinase assay was performed according to the method described by (Allen et al., 2007). Phosphorylation was detected by Western blot analysis using an anti-Thiophosphate ester antibody (ab92570; Abcam) and by the MS/MS analysis.

*In vivo* phosphorylation analysis was performed using a 2D western blot assay on nuclear extracts from 5 weeks old WT and *er/erl1/erl2* plants (Saleh et al., 2008). For isoelectrofocusing (IEF), nuclear proteins were prepared according to Kubala et al. (2015). The IEF was performed on the 7cm length gel strips with immobilized pH gradients 3-10 and 3-6 (BioRad). After IEF, the equilibration of immobilized pH gradient was performed according to Wojtyla et al. (2013). The SWI3B protein was detected by Western blotting using an anti SWI3B antibody (Sarnowski et al., 2002). The *in vivo* phosphorylation was identified based on the changes of SWI3B isoelectric point (*pI*) (Mayer et al., 2015).

### Accession Numbers

Microarray and MNase-seq data are available in the ArrayExpress database (www.ebi.ac.uk/arrayexpress) under E-MTAB-5595 and E-MTAB-5830 accession numbers, respectively.

## Supporting information

Supplemental data set 1

Supplemental data set 2

Supplemental data set 3

## Supplemental Data

**Supplemental Figure 1.** Comparative analysis of genes with altered expression in *er/erl1/erl2* and *ga1-3* mutants. A, Combinations of *erf* mutants have distinct effects on height of Arabidopsis plants. Error bars refer to SD,* = P < 0.05, Student’s t test. B, Genes showing altered expression in the *er/erl1/erl2* mutant plants. C, Venn diagrams indicating genes contrastingly up-regulated in *er/erl1/erl2* and down-regulated in *ga1-3* plants. D, Venn diagrams indicating genes contrastingly down-regulated in *er/erl1/erl2* and up-regulated in *ga1-3* plants. E, ERf proteins control the expression of genes down-regulated in *ga1-3,* which are either DELLA repressed or DELLA independent. F, ERf proteins control the expression of genes up-regulated in *ga1-3,* which are either DELLA activated or DELLA independent.

**Supplemental Figure 2.** The *er/erl1/erl2* mutant displays severely impaired response to exogenous GA treatment and slightly enhance *ga1-3* phenotypic traits. A, The 14 days old *er/erl1/erl2* plants treated with GA_4+7_ did not show rosette expansion indicating defects in GA response. Error bars refer to SD,* =P < 0.05, Student’s t test, n= 30 plants. B, Two-month-old GA_4+7_ treated *er/erl1/erl2* plants show accelerated flowering compared to mock treated control. Scale bar= 1cm. n= 30 plants. C, Two-months old *er/erl1/erl2* plants. Scale bar= 1cm. n= 30 plants. D, Three-weeks old *ga1-3* plants crossed with *erf* mutants in various combinations show mostly the phenotypic traits characteristic for *ga1-3*. Scale bar= 1 cm. E, The phenotypic traits of 5-weeks old plants carrying *erf* mutations crossed with *ga1-3* in various combinations grown in long day conditions. Scale bar= 1 cm.

**Supplemental Figure 3.** e*r/erl1/erl2* plants exhibit deficiency in gibberellin intermediates. Left panel: *er/erl1/erl2* mutant exhibits dramatically reduced level of GA_12_ and GA_24_ gibberellin intermediates (error bars-SD, P < 0.05, Student’s *t*-test, three biological and technical replicates were assayed). Right panel: schematic representation of alteration in GA biosynthesis pathway in *er/erl1/erl2* plants.

**Supplemental Figure 4.** 35S::ERECTA-GFP construct complements the *er-105* mutation.

**Supplemental Figure 5.** ERECTA protein is detected in various cell compartments including endosomes and accumulate in the nuclei periphery after BFA treatment. A, In cell pairs of stomata, ERECTA-GFP was detected in circles around the positions of nuclei. Scale bar=10µm. B, Accumulation of ERECTA protein in the BFA bodies after Brefeldin A treatment. Note the enhanced presence of BFA bodies after 40 min (mid column) and 90 min (right column) BFA treatment. Scale bar=10µm.

**Supplemental Figure 6.** The ERL1 and ERL2 proteins carry defined NLS in their kinase domains. A, The NLS prediction in the kinase domain of ERL1 and ERL2 proteins has been done using cNLS mapper (Kosugi et al., 2009a; Kosugi et al., 2009b). NLS score in range 5-7 means that protein is partially localized in the nucleus and cytoplasm. Bottom panel: The alignment of ERECTA, ERL1, and ERL2 protein sequences (part of kinase domains carrying NLS) using PRALINE indicates high amino-acid sequence conservation between analyzed proteins. Consistency is determined within range 1-10, where 1 means least conserved substitution and 10- the most conserved substitution (Simossis et al., 2005). B, Western blot analysis with anti-GFP antibody confirms nuclear localization of ERECTA protein which undergoes proteolytic processing. The samples were standardized by western blotting with anti H3 antibody. C, Schematic presentation of full length and deletion variants of ERECTA protein used for the localization study. ER – ERECTA protein with complete amino acids sequence; Δ ER – truncated ERECTA protein lacking kinase domain; KDER – the kinase domain of ERECTA protein.

**Supplemental Figure 7.** The kinase domain of ERECTA (KDER) has ability to complement the *er-105* leaf phenotypic traits. A, Rosette leaves of WT (upper), *er-105* (mid), and *er-105/*KDER-YFP (lower panel). Graphical alignment of corresponding leaves indicating partial complementation of *er* phenotypic traits by KDER. Scale bar= 1cm. B, Cauline leaves of WT (upper), *er-105* (mid), and *er-105/*KDER-YFP (lower panel). Graphical alignment of corresponding laves indicating partial complementation of *er* phenotypic traits by KDER. Scale bar= 1cm. C, The kinase domain of ERECTA cannot restore all (i.e., stem elongation) phenotypic traits of the *er-105* mutant line. Scale bar= 1cm.

**Supplemental Figure 8.** ERECTA protein enters to the nucleus and localizes in various sub-nuclear fractions.

**Supplemental Figure 9.** Negative controls for bimolecular fluorescence complementation assay. Negative controls for BiFC interaction analysis of ER, ERL1, ERL2 kinase domains fused to YFC and SWI3B fused to YFN, including the RFP channel. Scale bar 10 μ

**Supplemental Figure 10.** Human SWI3-type BAF155 co-precipitates with HER2 EGFR family membrane receptor from human cells nuclei.

**Supplemental Figure 11.** e*r/elr1/erl2* mutant plants show affected chromatin organization demonstrated as altered chromocenters number. Upper panel: exemplary pictures of WT and *er/erl1/erl2* nuclei. Lower panel: calculation of chromocenters (n=20 nuclei for each genotype).

**Supplemental Figure 12.** ERf proteins inactivation has a severe impact on genome-wide nucleosome positioning. A, Nucleosome changes identified in the *er/erl1/erl2* triple mutant plants. B, Genome-wide nucleosome distribution patterns surrounding the transcription start site (TSS). C, Nucleosome distribution patterns surrounding the TSS of GA-related genes showing altered expression in the *er/erl1/erl2* triple mutant plants. D, The alteration of nucleosomal structure on *PRE1*, *GID1a,* and *GID1b loci* misexpressed in the *er/erl1/erl2* triple mutant plants and targeted by the SWI/SNF CRC. Red boxes indicate nucleosome alterations. E, Confirmatory MNase-qPCR for selected genes with altered nucleosomes.

**Supplemental Figure 13.** Kinase domain of ERECTA phosphorylates SWI3B protein. A, Western blot with anti His6 antibody for detection of MBP-His6-KDER and His6-SWI3B proteins purified from bacteria. B, Western blot with anti-Thiophosphate ester antibody indicating no phosphorylation of SWI3B protein in the absence of KDER (negative control). C, Western blot with anti-Thiophosphate ester antibody (ab92570; Abcam) showing autophosphorylation of KDER in the absence of SWI3B protein. D, Identification of active phosphorylation sites in KDER by MS/MS analysis. E, Identification of active phosphorylation sites in SWI3B by MS/MS analysis. F, KDER phosphorylates SWI3B at the SWIRM and SANT domains.

**Supplemental Figure 14.** GA-deficient mutant lines with inactivated subunits of SWI/SNF complexes do not accumulate RGA DELLA protein. A, GA-deficient *swi3c* plants are unable to over accumulate RGA protein. B, GA-deficient *brm* plants are unable to over accumulate RGA protein.

**Supplemental Figure 15.** Negative controls for bimolecular fluorescence complementation assay.

Negative controls for BiFC interaction analysis of SWI3B, RGA, and RGL1 fused to YFC, and SWI3B fused to YFN, including the RFP channel. Scale bar 10 μm

**Supplemental Figure 16.** Proper SWI3B binding to *GID1a-c* promoter regions is abolished in the *er/erl1/erl2* whereas occurs in single *er-105, erl1* or *erl2* mutant lines. A, SWI3B targets promoter region of *GID1a* gene in five-weeks old plants WT, *er-105, erl1* or *erl2* but is abolished in triple *er/erl1/erl2* mutant plants (error bars refer to SD, P < 0.05, Student’s t test, three biological and technical replicates were used). B, SWI3B targets promoter region of *GID1b* gene in five-weeks old plants WT, *er-105, erl1* or *erl2* but is abolished in triple *er/erl1/erl2* mutant plants (error bars refer to SD, P < 0.05, Student’s t test, three biological and technical replicates were used). C, SWI3B targets promoter region of *GID1c* gene in five-weeks old plants WT, *er-105, erl1* or *erl2* but is abolished in triple *er/erl1/erl2* mutant plants (error bars refer to SD, P < 0.05, Student’s t test, three biological and technical replicates were used).

**Supplemental Figure 17.** Human SWI3-type BAF155 targets together with HER2 EGFR family membrane receptor *FBP1* and *BRCA1* genes *loci*. A, BAF155 subunit of human SWI/SNF complex binds *Fructose-1,6-Bisphosphatase locus* together with HER2 member of EGFR membrane receptor family. B, BAF155 subunit of human SWI/SNF complex binds *BRCA1 locus* together with HER2 member of EGFR membrane receptor family.

**Supplemental Table 1.** Genes classified to “Response to Gibberellin” GO term and showing down-regulated expression level in the *er/erl1/erl2* mutant.

**Supplemental Table 2.** Genes with up-regulated expression in *er/erl1/erl2* mutant plants classified to GO-terms of leaf epidermal and stomatal cell differentiation.

**Supplemental Table 3.** Functional analogies between arabidopsis ERf proteins and the human EGFR membrane receptors.

**Supplemental dataset 1.** Comparative analysis of transcript profiling and MNase-seq data.

**Sub-table 1.** Transcript profiling using ATH1 microarray analysis to identify genes down-regulated in *er/erl1/erl2* mutant line.

**Sub-table 2.** Transcript profiling using ATH1 microarray analysis to identify genes up-regulated in *er/erl1/erl2* mutant line.

**Sub-table 3.** GO analysis of genes down-regulated in *er/erl1/erl2* mutant line.

**Sub-table 4.** GO analysis of genes up-regulated in *er/erl1/erl2* mutant line.

**Sub-table 5.** Comparative transcript profiling analysis for genes down-regulated in *er/erl1/erl2* and *ga1-3* mutants lines.

**Sub-table 6.** GO analysis of *er/erl1/erl2* and *ga1-3* down-regulated genes.

**Sub-table 7.** Comparative transcript profiling analysis for genes up-regulated in *er/erl1/erl2* and *ga1-3* mutants lines.

**Sub-table 8.** GO analysis of genes up-regulated in *ga1-3* and *er/erl1/erl2*.

**Sub-table 9.** Genes with altered nucleosome positioning in promoter region -3000 to TSS.

**Sub-table 10.** GO analysis of genes with altered nucleosome positioning identified in promoter region -3000 to TSS.

**Sub-table 11.** Genes with altered nucleosome positioning identified in promoter region -3000 to TSS and down-regulated in *er/erl1/erl2* microarray.

**Sub-table 12.** Genes with altered nucleosome positioning identified in promoter region -3000 to TSS and up-regulated in *er/erl1/erl2* microarray.

**Sub-table 13.** GO analysis for genes with altered nucleosome positioning identified in promoter region -3000 to TSS and down-regulated in *er/erl1/erl2* microarray.

**Sub-table 14.** GO analysis for genes with altered nucleosome positioning identified in promoter region -3000 to TSS and up-regulated in *er/erl1/erl2* microarray.

**Supplemental dataset 2.** Comparison of ER, ERL1, and ERL2 protein sequences with highlighted important domains including NLS.

ClustalW was used to align ER, ERL1, and ERL2 sequences. Under conserved amino acid is asterisk mark, highly similar or similar amino-acids are marked :: and . respectively. LRR domain is marked as gray background, amino-acids crucial for EPFs or TMM interaction are indicated with bold. Transmembrane domain, juxtamembrane domain, and kinase domain are indicated with bold green, orange, and blue color fonts, respectively (Kosentka et al., 2017). The predicted NLS sequence is marked in a dotted line frame (Kosugi et al., 2009b).

**Supplemental Dataset 3.** Primers used in this work.

**Supplemental Movie 1. The ERECTA protein undergoes endocytosis.**

**Supplemental Movie 2. The ERECTA protein undergoes endocytosis.**

## Acknowledgements

We thank Csaba Koncz for critical comments during manuscript construction, Dorota Zugaj for assistance in plant cultivation, Iga Jancewicz for assistance in leptomycin B assay, Claus Schwechheimer for providing the anti-RGA antibody, Mohammad-Reza Hajirezaei for help with GA measurements and the Max Planck-Genome-Center Cologne (http://mpgc.mpipz.mpg.de/home/) for performing the transcript profiling and MNase-seq analysis described in this study.

## Authors Contributions

TJS, ES, and SJD planned experiments and wrote the manuscript

SK and PC participated in the planning of some experiments

TJS, ES, SK, PC, SS, SA, BH, JAS, and ARF analyzed the data

ES, PC, SS, SK, PO, JS, MZ, JMS, RD, AM, MS, BH, MC, KN, ATR, EB, RF, AK, MAD, SA, and TJS performed experiments

All authors read, edited, and approved the final manuscript

## Abbreviations and Acronyms

ERf: ERECTA family
ER: ERECTA
ERL1: ERECTA-LIKE 1
ERL2: ERECT-LIKE 2
LRR-RLKs: leucine-rich repeat receptor-like kinases
CRC: chromatin remodeling complex
SWI/SNF: Switch/Sucrose Nonfermenting
GID1: GIBBERELLIN INSENSITIVE DWARF 1
GA: gibberellin
PAC: Paclobutrazol
qRT-PCR: quantitative real-time PCR
BFA: Brefeldin A
NLS: nuclear localization signal
KDER: the kinase domain of ERECTA
TSS: transcription start site
EGFR: epidermal growth factor receptor

