## Supplemental data set 2 for "A Non-Canonical Function of Arabidopsis ERECTA Proteins in Gibberellin Signaling"

```
ER        MALFRDIVLLGFLFCLSL-------VATVTSEEGATLLEIKKSFKDVNNVLYDWTTSPSS
ERL1      ----MKEKMQRMVLSLAMVGFMVFGVASAMNNEGKALMAIKGSFSNLVNMLLDWDDVHNS
ERL2      --MRRIETMKGLFFCLGMVVFMLLGSVSPMNNEGKALMAIKASFSNVANMLLDWDDVHNH
                  :  :.:.*.:        .:  .:** :*: ** **.:: *:* **    .

ER        DYCVWRGVSCENVTFNVVALNLSDLNLDGEISPAIGDLKSLLSIDLRGNRLSGQIPDEIG
ERL1      DLCSWRGVFCDNVSYSVVSLNLSSLNLGGEISPAIGDLRNLQSIDLQGNKLAGQIPDEIG
ERL2      DFCSWRGVFCDNVSLNVVSLNLSNLNLGGEISSALGDLMNLQSIDLQGNKLGGQIPDEIG
          * * **** *:**: .**:****.***.**** *:*** .* ****:**:*.********

ER        DCSSLQNLDLSFNELSGDIPFSISKLKQLEQLILKNNQLIGPIPSTLSQIPNLKILDLAQ
ERL1      NCASLVYLDLSENLLYGDIPFSISKLKQLETLNLKNNQLTGPVPATLTQIPNLKRLDLAG
ERL2      NCVSLAYVDFSTNLLFGDIPFSISKLKQLEFLNLKNNQLTGPIPATLTQIPNLKTLDLAR
          :* **  :*:* * * ************** * ****** **:*:**:****** ****

ER        NKLSGEIPRLIYWNEVLQYLGLRGNNLVGNISPDLCQLTGLWYFDVRNNSLTGSIPETIG
ERL1      NHLTGEISRLLYWNEVLQYLGLRGNMLTGTLSSDMCQLTGLWYFDVRGNNLTGTIPESIG
ERL2      NQLTGEIPRLLYWNEVLQYLGLRGNMLTGTLSPDMCQLTGLWYFDVRGNNLTGTIPESIG
          *:*:*** **:************** *.*.:* *:************.*.***:***:**

ER        NCTAFQVLDLSYNQLTGEIPFDIGFLQVATLSLQGNQLSGKIPSVIGLMQALAVLDLSGN
ERL1      NCTSFQILDISYNQITGEIPYNIGFLQVATLSLQGNRLTGRIPEVIGLMQALAVLDLSDN
ERL2      NCTSFEILDVSYNQITGVIPYNIGFLQVATLSLQGNKLTGRIPEVIGLMQALAVLDLSDN
          ***:*::**:****:** **::**************:*:*:**.**************.*

ER        LLSGSIPPILGNLTFTEKLYLHSNKLTGSIPPELGNMSKLHYLELNDNHLTGHIPPELGK
ERL1      ELVGPIPPILGNLSFTGKLYLHGNMLTGPIPSELGNMSRLSYLQLNDNKLVGTIPPELGK
ERL2      ELTGPIPPILGNLSFTGKLYLHGNKLTGQIPPELGNMSRLSYLQLNDNELVGKIPPELGK
           * * ********:** *****.* *** ** ******:* **:****.*.* *******

ER        LTDLFDLNVANNDLEGPIPDHLSSCTNLNSLNVHGNKFSGTIPRAFQKLESMTYLNLSSN
ERL1      LEQLFELNLANNRLVGPIPSNISSCAALNQFNVHGNLLSGSIPLAFRNLGSLTYLNLSSN
ERL2      LEQLFELNLANNNLVGLIPSNISSCAALNQFNVHGNFLSGAVPLEFRNLGSLTYLNLSSN
          * :**:**:*** * * **.::***: **.:***** :**::*  *::* *:********

ER        NIKGPIPVELSRIGNLDTLDLSNNKINGIIPSSLGDLEHLLKMNLSRNHITGVVPGDFGN
ERL1      NFKGKIPVELGHIINLDKLDLSGNNFSGSIPLTLGDLEHLLILNLSRNHLSGQLPAEFGN
ERL2      SFKGKIPAELGHIINLDTLDLSGNNFSGSIPLTLGDLEHLLILNLSRNHLNGTLPAEFGN
          .:** **.**.:* ***.****.*::.* ** :******** :******:.* :*.:***

ER        LRSIMEIDLSNNDISGPIPEELNQLQNIILLRLENNNLTGNV-GSLANCLSLTVLNVSHN
ERL1      LRSIQMIDVSFNLLSGVIPTELGQLQNLNSLILNNNKLHGKIPDQLTNCFTLVNLNVSFN
ERL2      LRSIQIIDVSFNFLAGVIPTELGQLQNINSLILNNNKIHGKIPDQLTNCFSLANLNISFN
          ****  **:* * ::* ** **.****:  * *:**:: *:: ..*:**::*. **:*.*

ER        NLVGDIPKNNNFSRFSPDSFIGNPGLCGSWLNSPCHDSRRTVRVSISRAAILGIAIGGLV
ERL1      NLSGIVPPMKNFSRFAPASFVGNPYLCGNWVGSICGP-LPKSRV-FSRGALICIVLGVIT
ERL2      NLSGIIPPMKNFTRFSPASFFGNPFLCGNWVGSICGPSLPKSQV-FTRVAVICMVLGFIT
          ** * :*  :**:**:* **.*** ***.*:.* *     . :* ::* *:: :.:* :.

ER        ILLMVLIAACRPHNPPPFLDGSLDKPVTYSTPKLVILHMNMALHVYEDIMRMTENLSEKY
ERL1      LLCMIFLAVYKSMQQKKILQGSSKQAE--GLTKLVILHMDMAIHTFDDIMRVTENLNEKF
ERL2      LICMIFIAVYKSKQQKPVLKGSSKQPE--GSTKLVILHMDMAIHTFDDIMRVTENLDEKY
          :: *:::*. :  :   .*.** .:    .  *******:**:*.::****:****.**:

ER        IIGHGASSTVYKCVLKNCKPVAIKRLYSHNPQSMKQFETELEMLSSIKHRNLVSLQAYSL
ERL1      IIGYGASSTVYKCALKSSRPIAIKRLYNQYPHNLREFETELETIGSIRHRNIVSLHGYAL
ERL2      IIGYGASSTVYKCTSKTSRPIAIKRIYNQYPSNFREFETELETIGSIRHRNIVSLHGYAL
          ***:*********. *..:*:****:*.: * .:::****** :.**:***:***:.*:*
```

```
ER        SHLGSLLFYDYLENGSLWDLLHGPTKKKTLDWDTRLKIAYGAAQGLAYLHHDCSPRIIHR
ERL1      SPTGNLLFYDYMENGSLWDLLHGSLKKVKLDWETRLKIAVGAAQGLAYLHHDCTPRIIHR
ERL2      SPFGNLLFYDYMENGSLWDLLHGPGKKVKLDWETRLKIAVGAAQGLAYLHHDCTPRIIHR
          *  *.******:***********  ** .***:****** *************:******

ER        DVKSSNILLDKDLEARLTDFGIAKSLCVSKSHTSTYVMGTIGYIDPEYARTSRLTEKSDV
ERL1      DIKSSNILLDENFEAHLSDFGIAKSIPASKTHASTYVLGTIGYIDPEYARTSRINEKSDI
ERL2      DIKSSNILLDGNFEARLSDFGIAKSIPATKTYASTYVLGTIGYIDPEYARTSRLNEKSDI
          *:******** ::**:*:*******: .:*:::****:***************:.****:

ER        YSYGIVLLELLTRRKAVDDESNLHHLIMSKTGNNEVMEMADPDITSTCKDLGVVKKVFQL
ERL1      YSFGIVLLELLTGKKAVDNEANLHQLILSKADDNTVMEAVDPEVTVTCMDLGHIRKTFQL
ERL2      YSFGIVLLELLTGKKAVDNEANLHQMILSKADDNTVMEAVDAEVSVTCMDSGHIKKTFQL
          **:********* :****:*:***::*:**:.:* *** .* ::: ** * * ::*.***

ER        ALLCTKRQPNDRPTMHQVTRVLGSFMLSEQPPAATDTSATLAGSCYVDEYANLKTPHSVN
ERL1      ALLCTKRNPLERPTMLEVSRVLLSLVPSLQVAKKLPSLDHSTK----------KLQQENE
ERL2      ALLCTKRNPLERPTMQEVSRVLLSLVPSPPPK-KLPSPAKVQ-----------EGEERRE
          *******:* :**** :*:*** *:: *        :                :  .  :

ER        CSSMSASDAQLFLRFGQVISQNSE
ERL1      VRNPDAEASQWFVQFREVISKSSI
ERL2      SHSSDTTTPQWFVQFREDISKSSL
            . .:   * *::* : **:.*

Legend:
gray background - LRR domain (Kosentka et al., 2017 J. Exp. Bot.)
bold fonts & gray background – TMM&EPF interacting residues (Lin et al. 2017 Genes & Development)
green bold fonts – transmembrane domain (Kosentka et al., 2017 J. Exp. Bot.)
orange bold fonts – juxtamembrane domain (Kosentka et al., 2017 J. Exp. Bot.)
blue bold fonts – kinase domain (PFAM)
```

- predicted NLS (Kosugi et al., 2009, PNAS)
