## Supplemental tables for "A Non-Canonical Function of Arabidopsis ERECTA Proteins in Gibberellin Signaling"

**Supplemental Table 1.** Genes Classified to “Response to Gibberellin” GO Term and Showing Downregulated Expression Level in the *er/erl1/erl2* Mutant.

| **TAIR_ID** | **Name** |
| --- | --- |
| AT1G62660 | Glycosyl hydrolases family 32 protein(AT1G62660) |
| AT1G26960 | Homeobox protein 23(AtHB23) |
| AT1G12240 | Glycosyl hydrolases family 32 protein(ATBETAFRUCT4) |
| AT5G39860 | Basic helix-loop-helix (bHLH) DNA-binding family protein(PRE1) |
| AT2G04240 | RING/U-box superfamily protein(XERICO) |
| AT4G09460 | Myb domain protein 6(MYB6) |
| AT3G27810 | Myb domain protein 21(MYB21) |
| AT5G39760 | Homeobox protein 23(HB23) |
| AT5G67300 | Myb domain protein r1(MYBR1) |
| AT5G40350 | Myb domain protein 24(MYB24) |
| AT1G74650 | Myb domain protein 31(MYB31) |
| AT1G74660 | Mini zinc finger 1(MIF1) |
| AT1G07430 | Highly ABA-induced PP2C protein 2(HAI2) |
| AT2G46830 | Circadian clock associated 1(CCA1) |
| AT1G67030 | Zinc finger protein 6(ZFP6) |
| AT4G25420 | 2-oxoglutarate (2OG) and Fe(II)-dependent oxygenase superfamily protein(GA20ox1) |
| AT5G15160 | BANQUO 2(BNQ2) |
| AT5G11260 | Basic-leucine zipper (bZIP) transcription factor family protein(HY5) |
| AT5G59845 | Gibberellin-regulated family protein(AT5G59845) |
| AT4G26150 | Cytokinin-responsive gata factor 1(CGA1) |
| AT2G14900 | Gibberellin-regulated family protein(AT2G14900) |
| AT2G45900 | Phosphatidylinositol N-acetyglucosaminlytransferase subunit P-like protein(TRM13) |
| AT4G30270 | Xyloglucan endotransglucosylase/hydrolase 24(XTH24) |
| AT3G63010 | Alpha/beta-Hydrolases superfamily protein(GID1B) |
| AT1G66350 | RGA-like 1(RGL1) |
| AT4G25000 | Alpha-amylase-like protein(AMY1) |
| AT1G68360 | C2H2 and C2HC zinc fingers superfamily protein(AT1G68360) |

**Supplemental Table 2.** Genes with Up-regulated Expression in *er/erl1/erl2* Mutant Plants Classified to GO-terms of Leaf Epidermal and Stomatal Cell Differentiation.

| TAIR_ID | Name | Function |
| --- | --- | --- |
| AT1G02340 | basic helix-loop-helix (bHLH) DNA-binding superfamily protein(HFR1) | Encodes a light-inducible, nuclear bHLH protein involved in phytochrome signaling. Mutants exhibit a long-hypocotyl phenotype only under far-red light but not under red light and are defective in other phytochrome A-related responses. Mutants also show blue light response defects. HFR1 interacts with COP1, co-localizes to the nuclear speckles and is ubiquinated by COP1. |
| AT1G04110 | Subtilase family protein(SDD1) | stomatal complex morphogenesis |
| AT1G08810 | myb domain protein 60(MYB60) | putative transcription factor of the R2R3-MYB gene family. Transcript increases under conditions that promote stomatal opening (white and blue light, abi1-1 mutation) and decreases under conditions that trigger stomatal closure (ABA, desiccation, darkness), with the exception of elevated CO2. Expressed exclusively in guard cells of all tissues. It is required for light-induced opening of stomata. Mutant shows reduced stomatal aperture which helps to limit water loss during drought. |
| AT1G34245 | Putative membrane lipoprotein(EPF2) | stomatal development |
| AT1G80080 | Leucine-rich repeat (LRR) family protein(TMM) | stomatal complex morphogenesis |
| AT2G23760 | BEL1-like homeodomain 4(BLH4) | leaf morphogenesis |
| AT2G42870 | phy rapidly regulated 1(PAR1) | developmental process |
| AT2G46870 | AP2/B3-like transcriptional factor family protein(NGA1) | leaf and flower development |
| AT3G45780 | phototropin 1(PHOT1) | regulation of stomatal movement |
| AT3G49670 | Leucine-rich receptor-like protein kinase family protein(BAM2) | regulation of meristem growth and meristem structural organization |
| AT3G54720 | Peptidase M28 family protein(AMP1) | leaf vascular tissue pattern formation meristem, embryonic, root and flower development |
| AT5G53210 | basic helix-loop-helix (bHLH) DNA-binding superfamily protein(SPCH) | stomatal development |
| AT2G26580 | plant-specific transcription factor YABBY family protein(YAB5) | polarity specification of adaxial/abaxial axis, leaf development regulation, regulation of shoot apical meristem development, regulation of transcription, DNA-templated |

**Supplemental Table 3.** Functional Analogies Between Arabidopsis ERf Proteins and the Human EGFR Membrane Receptors.

| Family  members | | | ERECTA family proteins | | | EGFR/HER family proteins | | | | references |
| --- | --- | --- | --- | --- | --- | --- | --- | --- | --- | --- |
|  |  |  | ER | ERL1 | ERL2 | EGFR/HER1 | ErbB2/HER2 | ErbB3/HER3 | ErbB4/HER4 |  |
| localization | membrane | | yes | yes | yes | yes | yes | yes | yes | (Yarden and Sliwkowski, 2001; Shpak et al., 2004) |
|  | endosomes | | yes | ? | yes | yes | yes | yes | yes | (Yarden and Sliwkowski, 2001; Ho et al., 2016; Hsu and Hung, 2016) |
|  | nuclear | | yes | ?/Yes (kinase domain) | ?/Yes (kinase domain) | yes | yes | yes | yes | (Hsu and Hung, 2016) |
| Cleaved form | | | yes | ? | ? | yes | yes |  | yes | (Hsu and Hung, 2016) |
| Kinase activity | | | Serine/threonine kinase | Serine/threonine kinase | Serine/threonine kinase | Tyrosine kinase | Tyrosine kinase | Pseudokinase | Tyrosine kinase | (Karachaliou et al., 2016) |
| Directly targeted genes | | | GID1a, GID1b, GID1c | ? | ? | Aurora-A, B-myb, iNOS, cyclin D1, COX-2, c-Myc, BCRP | COX-2, | cyclin D1, | β-caseine | (Chen and Hung, 2015; Hsu and Hung, 2016) |
| Ligand binding | | | STOMAGEN, EPF2 | EPF1 | _ | EGF, TGF-α, HB-EGF, epiregulin, betacellulin, amphiregulin | _ | neuregulins | Neuregulins,  HB-EGF, epiregulin, betacellulin | (Hsu and Hung, 2016; Lee et al., 2015a; Tameshige et al., 2016; Wong, 2003) |
| dimerization | | homo | ? | yes | _ | yes | yes | _ | yes | (Yarden and Sliwkowski, 2001; Ho et al., 2016; Karachaliou et al., 2016) |
|  |  | hetero | yes | yes | yes | yes | yes | yes | yes |  |
| knockout plants/animals | | | phenotypical changes | no changes | no changes | Embryonic or perinatal lethality, in some strains viable with strong phenotypical changes | Embryonic lethality | Embryonic lethality | Embryonic lethality | (Miettinen et al., 1995; Torii et al., 1996; Casalini et al., 2004; Shpak et al., 2004) |
| Downstream signaling | | | MAPK cascade | MAPK cascade | MAPK cascade | MAPK cascade,  PI3K cascade, PLCγ/PKC, and JAK/STAT | MAPK cascade, PI3K cascade, PLCγ/PKC, and JAK/STAT | MAPK cascade, PI3K cascade, | PI3K cascade, MAPK cascade, PLCγ/PKC, and JAK/STAT | (Casalini et al., 2004; Hsu and Hung, 2016; Karachaliou et al., 2016; Tameshige et al., 2016) |
| Involvement in hormonal action | | | auxin, GA ethylene, | GA | GA | estrogen, androgen | androgen, | ? | estrogen | (Hsu and Hung, 2016; Uchida et al., 2012a, 2012b; Woodward and Bartel, 2005; Van Zanten et al., 2010) |
| Interaction with transcription factors | | | ? | ? | ? | STAT5,  STAT3,  E2F1 | STAT3 | ? | STAT5A, Eto-2, YAP | (Chen and Hung, 2015; Hsu and Hung, 2016; Hung et al., 2008) |
